# Resistance to decitabine and 5-azacytidine emerges from adaptive responses of the pyrimidine metabolism network

**DOI:** 10.1101/2020.02.20.958405

**Authors:** Xiaorong Gu, Rita Tohme, Benjamin Tomlinson, Metis Hasipek, Lisa Durkin, Caroline Schuerger, Asmaa M. Zidan, Tomas Radivoyevitch, Changjin Hong, Hetty Carraway, Betty Hamilton, Ronald Sobecks, Babal K. Jha, Eric D. Hsi, Jaroslaw Maciejewski, Yogen Saunthararajah

**Affiliations:** Department of Translational Hematology & Oncology Research, Taussig Cancer Institute, Cleveland Clinic, Cleveland, Ohio; Department of Hematology and Oncology, University Hospitals, Cleveland, Ohio; Department of Clinical Pathology, Tomsich Pathology Institute, Cleveland Clinic, Cleveland, Ohio; Department of Quantitative Health Sciences, Cleveland Clinic, Cleveland, Ohio; Department of Hematology and Oncology, Taussig Cancer Institute, Cleveland Clinic, Cleveland, Ohio

**Keywords:** 5-azacytidine, decitabine, DNMT1, myelodysplastic syndromes, leukemia, resistance, therapy

## Abstract

Mechanisms-of-resistance to decitabine and 5-azacytidine, mainstay treatments for myeloid malignancies, require investigation and countermeasures. Both are nucleoside analog pro-drugs processed by pyrimidine metabolism into a nucleotide analog that depletes the key epigenetic regulator DNA methyltranseferase 1 (DNMT1). We report here that DNMT1 protein, although substantially depleted (~50%) in patients’ bone marrows at response, rebounded at relapse, and explaining this, we found pyrimidine metabolism gene expression shifts averse to the processing of each pro-drug. The same metabolic shifts observed clinically were rapidly recapitulated in leukemia cells exposed to the pro-drugs in vitro. Pyrimidine metabolism is a network that senses and preserves nucleotide balances: Decitabine, a deoxycytidine analog, and 5-azacytidine, a cytidine analog, caused acute and distinct nucleotide imbalances, by off-target inhibition of thymidylate synthase and ribonucleotide reductase respectively. Resulting expression changes in key pyrimidine metabolism enzymes peaked 72-96 hours later. Continuous pro-drug exposure stabilized metabolic shifts generated acutely, preventing DNMT1-depletion and permitting exponential leukemia out-growth as soon as day 40. Although dampening to activity of the pro-drug initially applied, adaptive metabolic responses primed for activity of the other. Hence, in xenotransplant models of chemorefractory AML, alternating decitabine with 5-azacytidine, timed to exploit compensating metabolic shifts, and addition of an inhibitor of a catabolic enzyme induced by decitabine/5-azacytidine, extended DNMT1-depletion and time-to-distress by several months versus either pro-drug alone. In sum, resistance to decitabine and 5-azacytidine emerges from adaptive responses of the pyrimidine metabolism network; these responses can be anticipated and thus exploited.

**GRAPHICAL ABSTRACT:** 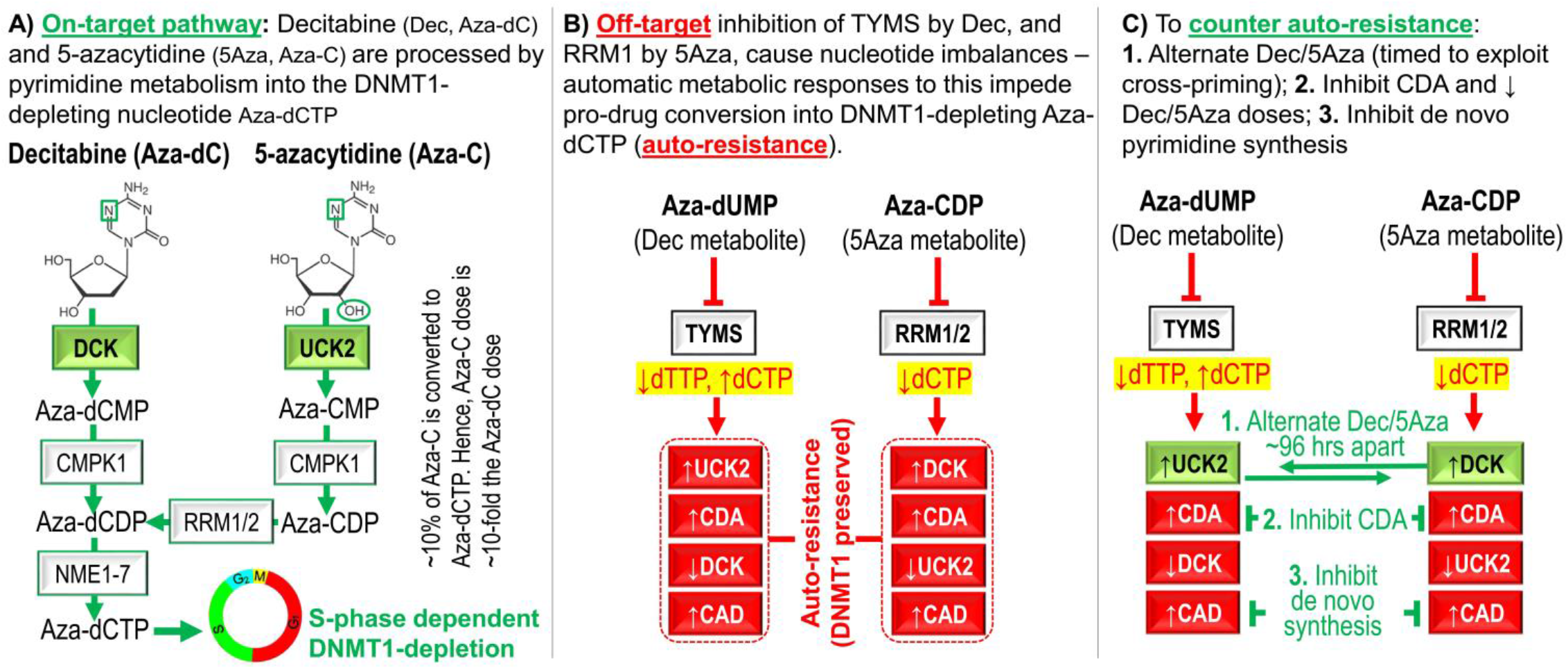

## INTRODUCTION

The deoxycytidine analog pro-drug decitabine and the cytidine analog pro-drug 5-azacytidine can increase life-spans of patients with myeloid malignancies, shown by randomized trials in patients with myelodysplastic syndromes (MDS) and acute myeloid leukemia (AML)(reviewed in^1^). Both pro-drugs are processed by pyrimidine metabolism into a deoxycytidine triphosphate (dCTP) analog, Aza-dCTP, that depletes the key epigenetic regulator DNA methyltransferase 1 (DNMT1) in dividing cells^2^. DNMT1-depletion terminates malignant self-replication but maintains normal hematopoietic stem cell self-replication^3–14^ – a vital therapeutic index when treating myeloid malignancies, since recovery by functional hematopoiesis is needed to reverse low blood counts, the cause of morbidity and death. Also clinically significant, the cell cycle exits induced by DNMT1-depletion do not require the p53 apoptosis axis, and can hence occur even in *TP53*-mutated, chemotherapy-resistant malignant cells (reviewed in^15^): Accordingly, decitabine or 5-azacytidine can benefit even patients with high risk, chemorefractory disease^1^. Nevertheless, only ~40% of treated patients benefit overall, and even in responders, relapse is typical. There is therefore a need to understand the mechanisms by which malignant cells resist decitabine or 5-azacytidine, and to use such knowledge to improve response rates and durations. An important piece of this puzzle could be how these pro-drugs are processed into DNMT1 targeting Aza-dCTP.

Decitabine and 5-azacytidine have an identical pyrimidine base modification - replacement of carbon at position 5 with nitrogen - but the sugar moiety is deoxyribose in decitabine and ribose in 5-azacytidine. This channels their metabolism differently, with critical roles for the following enzymes: deoxycytidine kinase (DCK), uridine cytidine kinase 2 (UCK2), cytidine deaminase (CDA), and carbamoyl-phosphate synthetase (CAD). The initial, rate-limiting step in the processing of decitabine toward Aza-dCTP is its phosphorylation by DCK^16,17^. *DCK*-null AML cells thus resisted decitabine, even at a concentration of 360μM^18^, and sensitivity was restored by transfection with an expression vector for DCK^16,19^. The initial phosphorylation of 5-azacytidine on the other hand is by UCK2^20,21^. Thus, AML cell lines resistant to >50μM of 5-azacytidine contained inactivating mutations in *UCK2*^22^, and sensitivity was restored by transfection with an expression vector for UCK2^22^. Despite such in vitro data, contributions of altered DCK and/or UCK2 to clinical relapse has been minimally investigated: one study of 14 decitabine-treated patients measured DCK expression in peripheral blood or bone marrow at relapse *vs* diagnosis, with inconclusive results^23^; another study of 8 decitabine-treated patients did find that DCK expression was significantly decreased at relapse^24^.

CDA is another pyrimidine metabolism enzyme shown to mediate resistance to decitabine or 5-azacytidine in vitro: CDA rapidly catabolizes both pro-drugs into uridine counterparts that do not deplete DNMT1^25^ and that instead cause off-target anti-metabolite effects, e.g., by misincorporating into DNA^26^. Thus, introduction of expression vectors for CDA into malignant cells conferred decitabine-resistance^27,28^. High CDA expression in tissues such as the liver underlies the brief in vivo plasma half-lives of decitabine and 5-azacytidine of ~15 minutes *vs* 9-16 hours in vitro at 37°C^29,30^. In a pre-clinical in vivo model, CDA-rich tissue micro-environments (e.g., liver) provided sanctuary to AML cells from decitabine^31^. In clinical analyses, higher CDA expression in males appeared to contribute to inferior responses of their MDS to decitabine or 5-azacytidine^32–34^. Nevertheless, as for DCK and UCK2, a contribution of CDA to clinical relapse is neither established nor addressed by current clinical practice.

CAD is the first enzyme in the de novo pathway that synthesizes dCTP from glutamine and aspartate: de novo synthesized dCTP can compete with Aza-dCTP for incorporation into DNA, and CAD upregulation has been implicated in resistance to 5-azacytidine in vitro^20,35,36^. Again, however, a contribution of CAD to clinical resistance/relapse has not been examined. Altogether therefore, DCK, UCK2, CDA and CAD expression changes are known to mediate resistance to decitabine or 5-azacytidine in vitro, but there is little information and no countermeasures for their individual or collective contributions to clinical resistance. Here, upon a first serial analyses of DNMT1 levels in patients’ bone marrows on clinical decitabine or 5-azacytidine therapy, we found that this target was not being engaged at clinical relapse and showing why, bone marrows at relapse exhibited shifts in DCK, UCK2, CDA and CAD expression in directions adverse to pro-drug conversion to Aza-dCTP. Pyrimidine metabolism is a network that senses and regulates nucleotide levels^37^, and we found that decitabine and 5-azacytidine cause distinct nucleotide imbalances, that in turn trigger specific, adaptive changes in expression of key pyrimidine metabolism enzymes. The consistency and predictability of pyrimidine metabolism network responses to decitabine or 5-azacytidine perturbations enabled their anticipation and exploitation to instead enhance pro-drug effects: simple, practical treatment modifications, evaluated in pre-clinical in vivo models of aggressive chemo-refractory AML, preserved the favorable therapeutic index of non-cytotoxic DNMT1-depletion and markedly improved efficacy.

## METHODS

### Study approvals

Bone marrow samples for research were obtained from patients with AML with written informed consent on a study protocol approved by the Cleveland Clinic Institutional Review Board (Cleveland, Ohio). Murine experiments were approved by the Cleveland Clinic Institutional Animal Care and Use Committee (Cleveland, Ohio).

### Detailed methods in Supplement

## RESULTS

### DNMT1 is not depleted at clinical relapse or with in vitro resistance

We measured DNMT1 protein levels in patients’ bone marrows before and during therapy with decitabine or 5-azacytidine (39 serial bone marrow samples from 13 patients, median treatment duration 372 days, range 170-1391). Serial bone marrow biopsies from the same patient were cut onto the same glass slide and stained simultaneously to facilitate time-course comparison of DNMT1 protein levels quantified by immunohistochemistry and ImageIQ imaging and software (**Figure 1A**). At time-of-response, DNMT1 protein was markedly and significantly decreased by ~50% compared to pre-treatment (**Figure 1A**). At the time-of-relapse on-therapy, however, DNMT1 protein levels had rebounded to levels comparable to or exceeding pre-treatment levels (**Figure 1A**).

**Figure 1.**
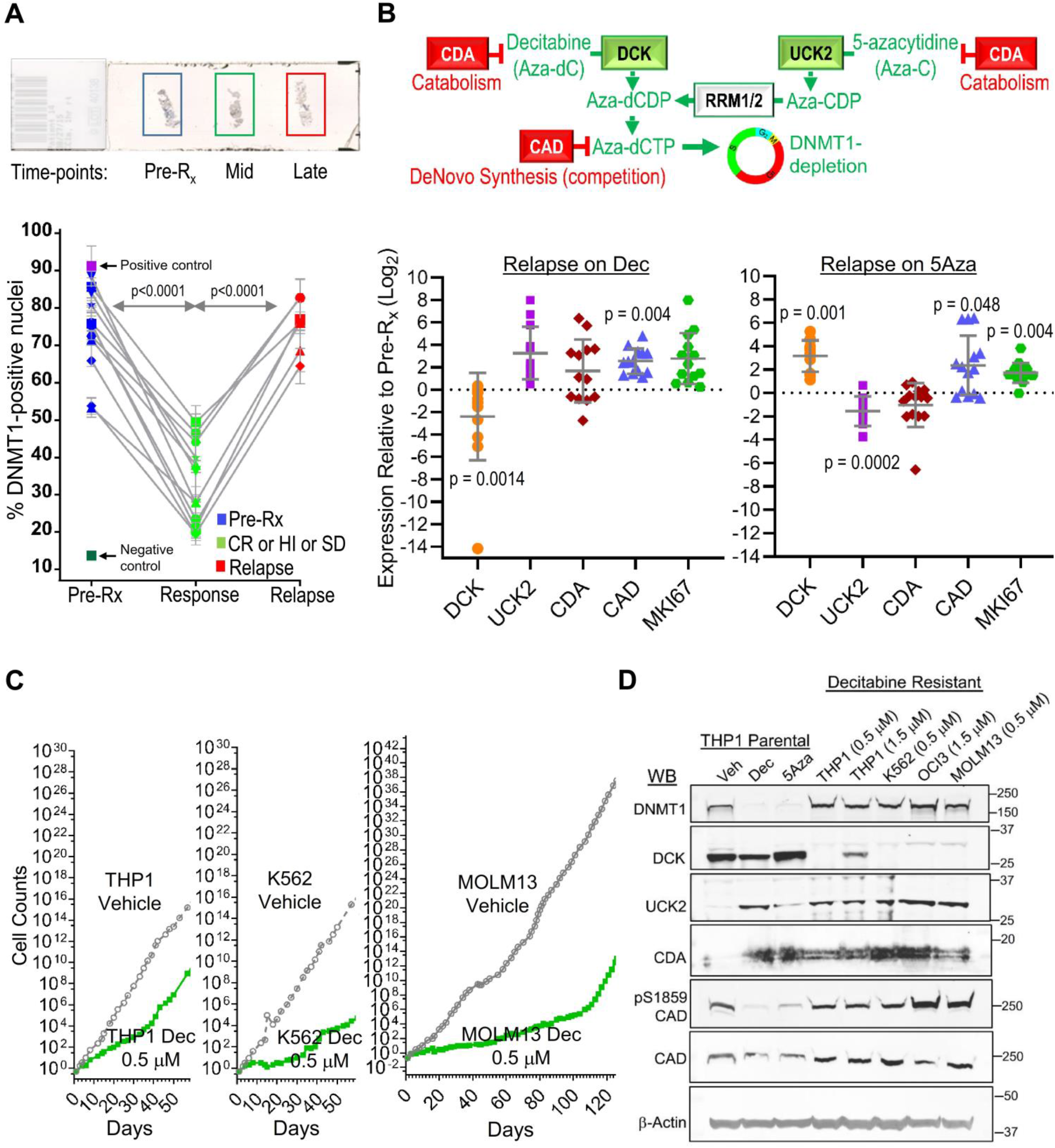
DNMT1 is not depleted at clinical relapse or with in vitro resistance. **A) Decitabine (Dec) or 5-azacytidine (5Aza) therapy decreased bone marrow DNMT1 at response (green) but DNMT1 rebounded to pre-treatment levels (dark blue) at relapse (red).** Serial bone marrow biopsies from the same patent were cut onto the same slide, stained for DNMT1, and the number of DNMT1-positive nuclei was quantified objectively using ImageIQ software, in 13 individual patients and positive/negative controls (tissue blocks of HCT116 wild-type and *DNMT1*-knockout cells respectively). D = days of therapy. Pre-R_x_ = pre-treatment; HI = hematologic improvement; CR = complete remission; SD = stable disease; Rel. = relapse. Mean±SD of ≥3 image segments (cellular regions) per sample; p-value paired t-test, 2-sided. **B) Expression of key pyrimidine metabolism enzymes at relapse/progression on Dec *vs* baseline.** Cartoon shows key enzymes favoring (green) or impeding (red) Dec or 5Aza conversion into the DNMT1-depleting Aza-dCTP. Bone marrow cells aspirated pre-treatment and at relapse/progression on Dec (13 patients, median duration of therapy 175 days, range 97-922) or 5Aza (14 patients, median duration of therapy 433 days, range 61-1155) were analyzed by QRT-PCR. Mean±SD technical replicatesx3. Paired t-test, 2-sided. **C) Time to emergence of AML cells exponentially proliferating in presence of Dec.** Cell counts by automated counter. **D) Expression by Western blot of key pyrimidine metabolism enzymes in Dec-resistant AML cells** (THP1, K562, OCI-AML3 and MOLM13), and in parental THP1 AML cells treated with vehicle, Dec or 5-azacytidine (5Aza). Western blots.

Since the pyrimidine metabolism enzymes DCK, UCK2, CDA and CAD are well-documented to mediate DNMT1-depleting capacity of decitabine and 5-azacytidine in vitro, we used quantitative polymerase chain reaction (QRT-PCR) to measure their expression in MDS patients’ bone marrows pre-treatment and at relapse on-therapy with decitabine (n=13, median treatment-duration 175 days, range 97-922) or 5-azacytidine (n=14, median treatment-duration 433 days, range 61-1155) (**Figure 1B**). At relapse on decitabine, expression of UCK2 and CDA increased ~8-fold and ~3-fold respectively, while DCK was decreased by ~50% *vs* pre-treatment levels (**Figure 1B**). By contrast, at relapse on 5-azacytidine, DCK expression increased ~8-fold while UCK2 and CDA expression decreased by ~50% (**Figure 1B**). Observed at relapse on both decitabine or 5-azacytidine, but not in all patients, were ~8-fold increases in expression of de novo pyrimidine synthesis enzyme CAD (**Figure 1B**). The proliferation marker MKI67 was increased at relapse in almost all the patients, consistent with active progression of disease (**Figure 1B**).

We then evaluated if AML cells resist clinically relevant concentrations of decitabine in vitro in a similar way. AML cells (THP1, K562, OCI-AML3, MOLM13) were cultured in the presence of decitabine 0.5 – 1.5μM. AML cells that proliferated exponentially in the presence of decitabine emerged as early as 40 days after the first addition of decitabine (**Figure 1C**). As per clinical relapse, DNMT1 was not depleted from the decitabine-resistant AML cells despite the presence of decitabine (**Figure 1D**). UCK2 and CDA protein levels were markedly elevated in decitabine-resistant *vs* vehicle-treated parental AML cells (**Figure 1D**). Also upregulated was CAD, by total protein levels and by S1856 phosphorylation, a post-translational modification linked with its functional activation (**Figure 1D**). In contrast, DCK protein levels were suppressed (**Figure 1D**). In short, the in vitro resistance resembled the clinical resistance, with preserved DNMT1 and reconfigured pyrimidine metabolism.

### Decitabine and 5-azacytidine cause acute nucleotide imbalances and metabolic shifts

Pyrimidine metabolism is a network that senses and regulates nucleotide levels^37^. Therefore, we examined whether decitabine and 5-azacytidine cause nucleotide imbalances. AML cells (MOLM13, OCI-AML3, THP1) were treated with a single dose of vehicle, natural deoxycytidine 0.5 μM, decitabine 0.5 μM, natural cytidine 5 μM, or 5-azacytidine 5 μM in vitro, and effects on nucleotide levels and pyrimidine metabolism gene expression were measured 24 to 72 hours later (**Figure 2A**). Vehicle, natural deoxycytidine and cytidine did not impact proliferation of the AML cells, while a single treatment with either decitabine or 5-azacytidine significantly decreased AML cell proliferation as expected (**Figure 2B**). Decitabine decreased dTTP and increased dCTP at 24 hours (**Figure 2C**). By contrast, 5-azacytidine decreased dCTP (**Figure 2C**). Decitabine consistently and significantly increased UCK2 and CDA mRNA (>2-fold), while 5-azacytidine consistently increased DCK and CDA mRNA (**Figure 2D**). Protein levels tracked the mRNA expression changes: decitabine upregulated UCK2 and downregulated DCK, with changes peaking between 48-96 hours (**Figure 2E, S1**). 5-azacytidine did the opposite: upregulated DCK and downregulated UCK2. Both pro-drugs upregulated CDA (**Figure 2E**). Thymidylate synthase (TYMS) is the major mediator of deoxythymidine triphosphate (dTTP) production^38–40^. TYMS was downregulated by both pro-drugs, but to a noticeably greater extent by decitabine than 5-azacytidine (**Figure 2E, S1**). 5-azacytidine, but not decitabine, decreased levels of the ribonucleotide reductase sub-unit RRM1 (**Figure 2E**). 5-azacytidine inconsistenly decreased levels of the ribonucleotide reductase sub-unit RRM2A (**Figure S1**). Neither pro-drug changed total CAD or CTPS1 levels (CTPS1 executes a late step in de novo pyrimidine synthesis) (**Figure 2E, S1**). Both pro-drugs did, however, decrease phosphorylation of CAD at serine 1856 (S1856)(**Figure 2E**) – a modification linked with functional upregulation of CAD-mediated de novo pyrimidine synthesis - thus, the pro-drugs acutely downregulated CAD function. Both decitabine (0.25μM) and 5-azacytidine (2.5μM) depleted DNMT1 as expected (**Figure 2E, S1**).

**Figure 2.**
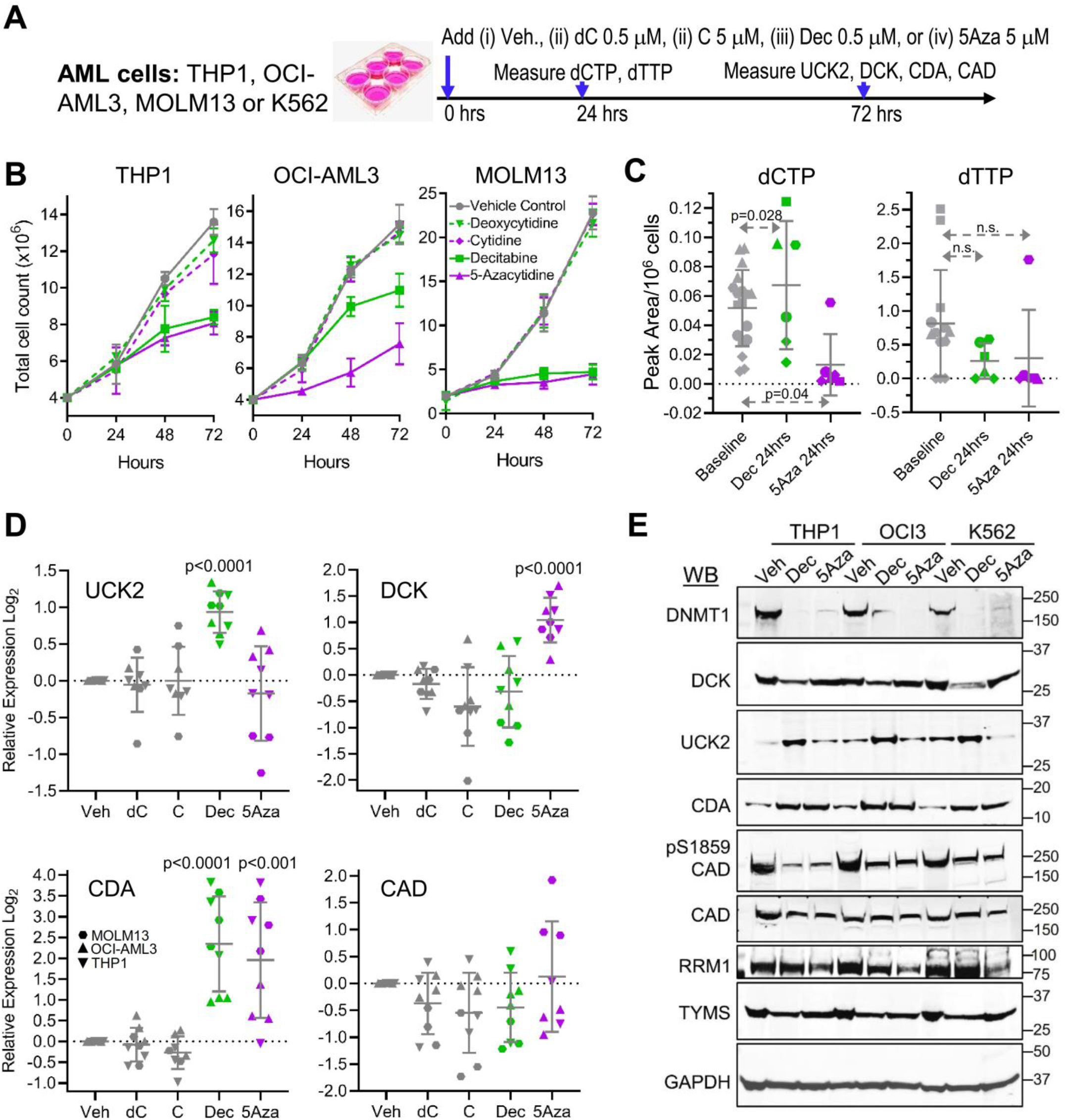
Dec or 5Aza, but not natural cytosine (C) or deoxycytosine (dC), rapidly altered expression of key pyrimidine metabolism enzymes UCK2, DCK and CDA in AML cells. **A) Experiment schema.** Vehicle, dC 0.5 μM, C 5 μM, Dec 0.5 μM, or 5Aza 5 μM were added once to the cells at 0 hours. Cell counts by automated counter**. B) Cell counts**. Means±SD for 3 independent biological replicates for each cell line. **C) Dec and 5Aza have opposite effects on dCTP levels.** Measured by LCMS/MS 24 hrs after addition of Dec or 5Aza. Analyses of 2 or more independent nucleotide extractions from 3 different AML cells lines. Means±SD; p-values paired t-test, 1-sided. **D) Gene expression 72 hours after Dec or 5Aza.** Gene expression by QRT-PCR, relative to average expression in vehicle-treated controls. Means±SD for 3 independent biological replicates in each of 3 AML cell lines; p-values unpaired t-test *vs* vehicle, 2-sided. **E) Western blot 72 hours after Dec or 5Aza.** AML cells THP1, OCI-AML3 and K562. Western blots were reproduced in three independent biological replicates.

### DCK is important for maintaining dCTP and UCK2 for maintaining dTTP levels

To examine DCK and UCK2 roles in dCTP and/or dTTP maintenance, we measured nucleotide levels in *DCK knock-out (KO)*, *UCK2* KO and wild-type control leukemia cells (HAP1). Knock-out of *DCK* significantly decreased dCTP but not dTTP (**Figure 3A, B**). Thus, DCK is important to dCTP maintenance, consistent with DCK upregulation as a response to dCTP suppression by 5-azacytidine (**Figure 2**). Knock-out of *UCK2* significantly decreased dTTP (**Figure 3A, B**). Thus, UCK2 is important to dTTP maintenance, consistent with UCK2 upregulation as a response to dTTP suppression by decitabine (**Figure 2**).

**Figure 3.**
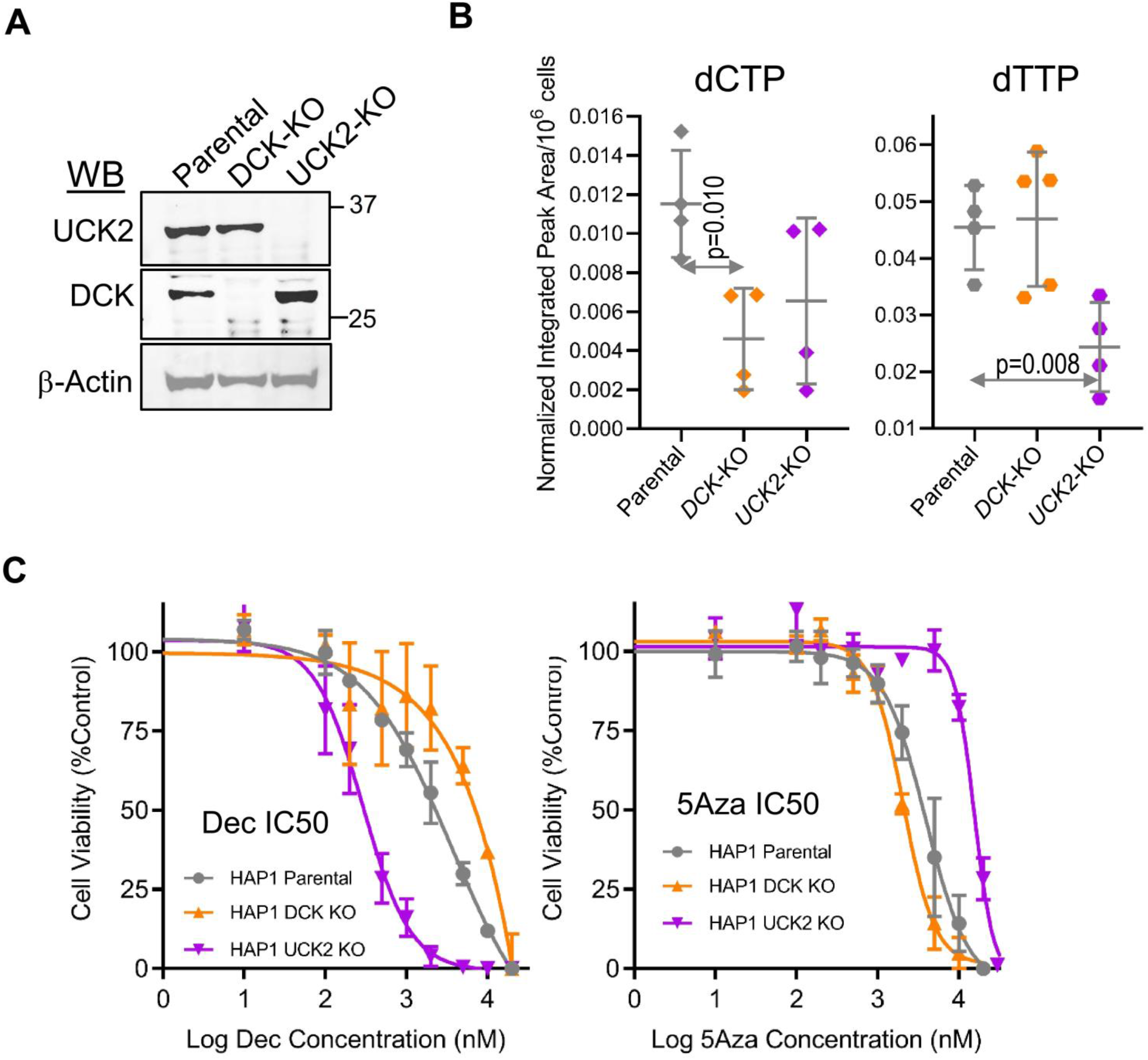
DCK is important for maintaining dCTP and UCK2 for maintaining dTTP levels, and DCK and UCK2 knock-outs do not create cross-resistance to both Dec and 5Aza. **A) Confirming DCK and UCK2 knockout (KO) by Western blot**. HAP1 leukemia cells, KO by CRISPR-Cas9. **B) *DCK*-KO lowers dCTP and *UCK2*-KO lowers dTTP.** Analysis of independent nucleotide extractions. Means±SD; p-values unpaired t-test, 2-sided. **C) Concentrations of Dec (left-panel) or 5Aza (right-panel) needed for 50% growth inhibition of parental, DCK-KO and UCK2-KO cells.** Means±SD of 3 independent biological replicates.

We also examined sensitivity of *DCK*- and *UCK2*-KO cells to decitabine or 5-azacytidine mediated growth inhibition. *DCK*-KO leukemia cells were relatively resistant to decitabine (concentrations for 50% growth inhibition [GI50] 12μM *vs* 3μM for wild-type), but more sensitive to 5-azacytidine (GI50 2μM *vs* 4μM for wild-type)(**Figure 3C**). *UCK2*-KO leukemia cells were relatively resistant to 5-azacytidine (GI50 15μM *vs* 4μM for wild-type), but more sensitive to decitabine (GI50 0.1μM *vs* 3μM for wild-type)(**Figure 3C**).

### Resistance countermeasures evaluated in vivo

We then examined solutions to resistance in a patient-derived xenotransplant (PDX) model of chemorefractory AML (summarized in **Table 1**):

**Table 1:**
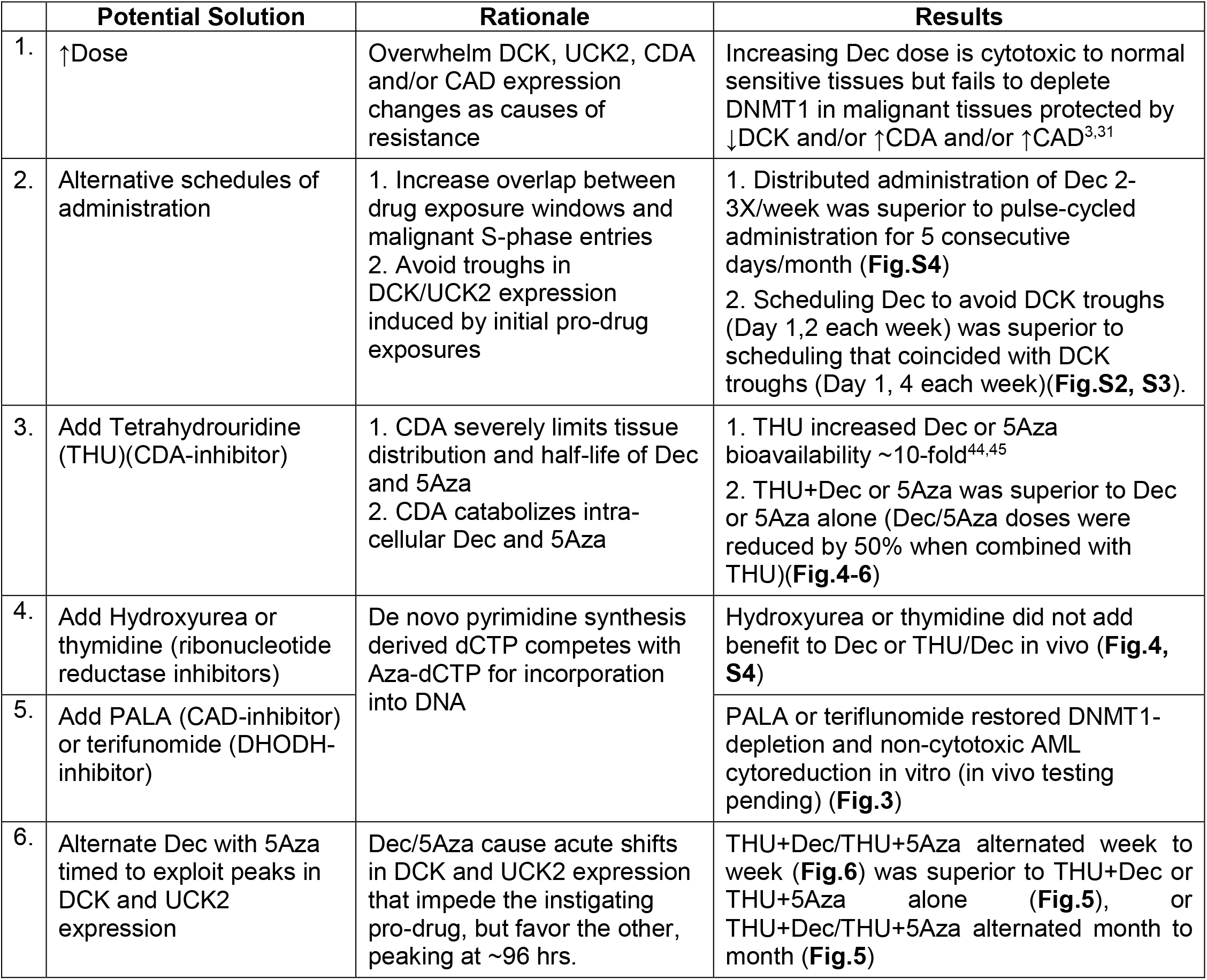
Solutions for resistance to decitabine and 5-azacytidine evaluated in vivo

|  | Potential Solution | Rationale | Results |
| --- | --- | --- | --- |
| 1. | ↑Dose | Overwhelm DCK, UCK2, CDA and/or CAD expression changes as causes of resistance | Increasing Dec dose is cytotoxic to normal sensitive tissues but fails to deplete DNMT1 in malignant tissues protected by ↓DCK and/or ↑CDA and/or ↑CAD<sup>3,31</sup> |
| 2. | Alternative schedules of administration | 1. Increase overlap between drug exposure windows and malignant S-phase entries<br>2. Avoid troughs in DCK/UCK2 expression induced by initial pro-drug exposures | 1. Distributed administration of Dec 2-3X/week was superior to pulse-cycled administration for 5 consecutive days/month (**Fig.S4**)<br>2. Scheduling Dec to avoid DCK troughs (Day 1,2 each week) was superior to scheduling that coincided with DCK troughs (Day 1, 4 each week)(**Fig.S2, S3**). |
| 3. | Add Tetrahydrouridine (THU)(CDA-inhibitor) | 1. CDA severely limits tissue distribution and half-life of Dec and 5Aza<br>2. CDA catabolizes intra-cellular Dec and 5Aza | 1. THU increased Dec or 5Aza bioavailability ~10-fold<sup>44,45</sup><br>2. THU+Dec or 5Aza was superior to Dec or 5Aza alone (Dec/5Aza doses were reduced by 50% when combined with THU)(**Fig.4-6**) |
| 4. | Add Hydroxyurea or thymidine (ribonucleotide reductase inhibitors) | De novo pyrimidine synthesis derived dCTP competes with Aza-dCTP for incorporation into DNA | Hydroxyurea or thymidine did not add benefit to Dec or THU/Dec in vivo (**Fig.4, S4**) |
| 5. | Add PALA (CAD-inhibitor) or terifunomide (DHODH-inhibitor) |  | PALA or terifunomide restored DNMT1-depletion and non-cytotoxic AML cyto-reduction in vitro (in vivo testing pending) (**Fig.3**) |
| 6. | Alternate Dec with 5Aza timed to exploit peaks in DCK and UCK2 expression | Dec/5Aza cause acute shifts in DCK and UCK2 expression that impede the instigating pro-drug, but favor the other, peaking at ~96 hrs. | THU+Dec/THU+5Aza alternated week to week (**Fig.6**) was superior to THU+Dec or THU+5Aza alone (**Fig.5**), or THU+Dec/THU+5Aza alternated month to month (**Fig.5**) |

#### Schedule decitabine administration to avoid DCK troughs

Immune-deficient mice were tail-vein innoculated with 1 million human AML cells each obtained from a patient with AML that was refractory/relapsed to decitabine then cytarabine. On Day 9 after innoculation, mice were randomised to treatment with **(i)** vehicle; **(ii)** decitabine timed to avoid DCK troughs (Day 1 and Day 2 each week – decitabine-Day1/2); or **(iii)** decitabine timed to coincide with DCK troughs (Day 1 and Day 4 each week – decitabine-Day1/4) (**Figure S2A**). Vehicle-treated mice showed distress on Day 45, at which point all mice were euthanized or sacrificed for analyses. The bone marrows of vehicle and also decitabine-Day1/4-treated mice, were replaced by AML cells by light microscope examination, but normal murine myelopoiesis was evident with decitabine-Day1/2 treatment (**Figure S2B**). This impression was confirmed by flow cytometry analyses of the bone marrows: human CD45+ (hCD45+) cells were ~92% with PBS, ~63% with decitabine-Day1/4 and ~26% with decitabine-Day1/2 treatment (**Figure S2C**). Spleens were enlarged and had effaced histology with vehicle or decitabine-Day1/4 treatment but had mostly preserved histology with decitabine-Day1/2 (**Figure S2D,E**). Spleen weights as another measure of AML burden were >5-fold greater with vehicle *vs* decitabine-Day1/4 but were lowest with decitabine-Day1/2 treatment (**Figure S2F**).

These two schedules of decitabine administration were compared again but with waiting for signs of distress rather than sacrifice at day 45 (**Figure S3A**). Median survival (time-to-distress) was significantly greater with decitabine-Day1/2 (75 days) *vs* decitabine-Day1/4 (60 days) or vehicle (40 days) (**Figure S3B**). Spleen AML burden was again lowest with decitabine-Day1/2 (~0.13 g) *vs* decitabine-Day1/4 (~0.59 g) or vehicle (~0.61 g)(**Figure S3C,D**). Thus, scheduling decitabine administration to avoid DCK troughs was superior to scheduling that coincided with these troughs.

#### Inhibit CDA / Inhibit ribonucleotide reductase in the de novo pyrimidine synthesis pathway / Schedule decitabine administrations to increase overlaps between drug exposure windows and malignant cell S-phase entries

CDA can be inhibited by THU, while de novo pyrimidine synthesis can be inhibited with deoxythymidine (dT) that inhibits ribonucleotide reductase in this pathway^41^. NSG mice tail-vein innoculated with 1 million AML cells each were randomised to **(i)** vehicle; **(ii)** THU+dT, **(iii)** decitabine; **(iv)** THU+decitabine; or **(v)** THU+dT+decitabine (**Figure 4A**). PBS and THU/dT-treated mice developed signs of distress, and were euthanized, on day 42. Mice receiving other treatments were sacrificed 3 weeks later (day 63) to increase chances of seeing differences in AML burden between these treatments (**Figure 4A**). Visual inspection of femoral bones of vehicle or THU/dT treated mice showed replacement of reddish functional hematopoiesis with whitish leukemia (**Figure 4B**), whereas femurs from THU/decitabine or THU/dT/decitabine-treated mice retained a normal reddish appearance (**Figure 4B**). Accordingly, flow cytometry demonstrated replacement by human AML cells in PBS and THU/dT-treated mice (>90% hCD45+), that was improved to some extent by decitabine alone (~85% hCD45+) but and even greater extent by either THU/decitabine (~35% hCD45+) or THU/dT/decitabine (~42% hCD45+)(**Figure 4C**)(dT did not add further to the benefit from THU). Murine hematopoiesis was completely suppressed with PBS and THU/dT (0% murine Cd45+), almost completely suppressed with decitabine-alone (~5% mCd45+) but similarly preserved with THU/decitabine (~40% Cd45+) or THU/dT/decitabine treatment (~27% Cd45+)(**Figure 4C**). Hemoglobin and platelet levels were most suppressed, and white cell (peripheral leukemia) counts most elevated, with vehicle or THU/dT treatment, but only mildly to moderately suppressed with any of the decitabine containing regimens (**Figure 4D**). Spleen weights as a measure of AML burden were ~4-fold greater with vehicle or THU/dT *vs* decitabine-alone treatment, and lowest with THU/decitabine- or THU/dT/decitabine (**Figure 4E**). Spleen histology confirmed replacement by AML cells (with necrotic areas) with vehicle or THU/dT (**Figure 4E**), AML infiltration with decitabine-alone, but normal-appearance with THU/decitabine or THU/dT/decitabine treatment (**Figure 4E**). We also evaluated the use of hydroxyurea to inhibit ribonucleotide reductase: hydroxyurea 100 mg/kg IP was administered on Day 1 before THU/decitabine on days 2 and 3 - hydroxyurea addition did not add benefit (**Figure S4**). Thus, using THU to inhibit CDA augmented decitabine anti-AML activity but incorporation of dT or hydroxyurea to inhibit de novo pyrimidine synthesis did not.

**Figure 4.**
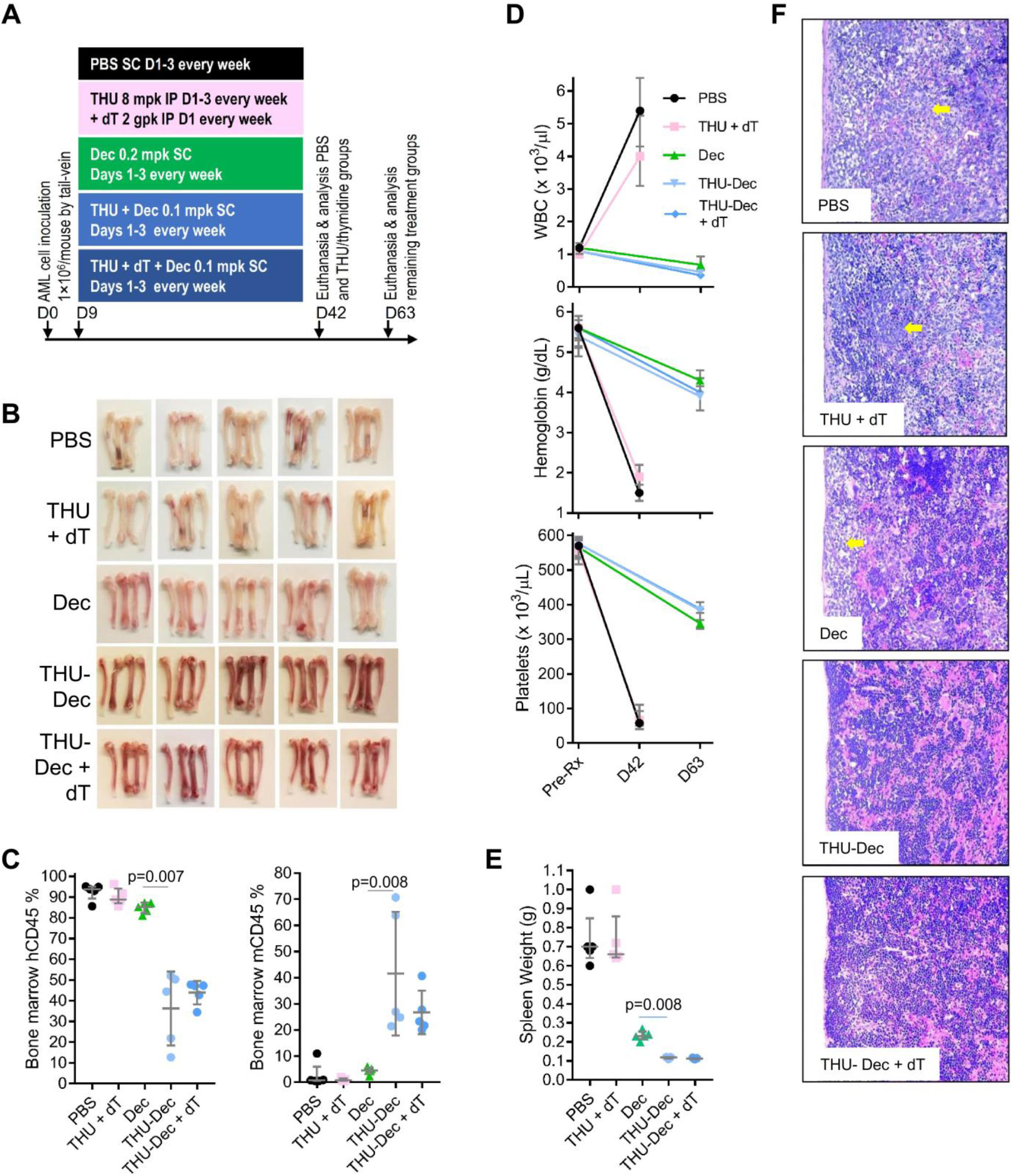
Impact of inhibiting CDA and/or de novo pyrimidine synthesis on Dec efficacy. NSG mice were tail-vein inoculated with patient-derived AML cells (1×10^6^cells/mouse) and randomized to (i) PBS vehicle control; (ii) CDA-inhibition by intra-peritoneal (IP) tetrahydrouridine (THU) and de novo pyrimidine synthesis inhibition by IP thymidine (dT); (iii) Dec; (iv) THU and Dec; (v) THU, Dec and dT (n=5/group). PBS, THU+dT mice were euthanized for distress on D42, and other mice were sacrificed for analysis on D63. **A) Experiment schema**; **B) Femoral bones** (from 2 of 5 mice/group). White = leukemia replacement, reddish = functional hematopoiesis. **C) Bone marrow human (hCD45) and murine (mCd45) myelopoiesis content.** Flow-cytometry. Median±IQR. p-value Mann-Whitney test 2-sided. **D) Blood counts before treatment and at euthanasia/sacrifice**. Measured by Hemavet. Median±IQR. **E) Spleen AML burden as measured by** s**pleen weights at euthanasia/sacrifice**. Median±IQR. p-value Mann-Whitney test 2-sided. **F) Spleen histology**. Hematoxylin-Eosin stain of paraffin-embedded sections. Magnification 400X. Leica DMR microscope.

DNMT1-depletion by decitabine or 5-azacytidine is S-phase dependent, suggesting frequent, distributed administration, to increase chances of overlap between malignant S-phase entries and drug exposures, could be better than historical pulse-cycled administration. Accordingly, bone marrow AML burden was lowest with frequent, distributed administration of THU/decitabine 2X/week (Day 1,2) compared to pulse-cycled administration of THU/decitabine for 5 days every 4 weeks (**Figure S4**).

#### Alternate THU/decitabine with THU/5-azacytidine (to exploit priming by Dec for 5Aza and vice-versa)

We compared head-to-head THU/decitabine *vs* THU/5-azacytidine and found no differences in efficacy between these two treatments (**Figure 5, S5**). Then, since decitabine appears to cross-prime for 5-azacytidine activity by upregulating UCK2, while 5-azacytidine cross-primes for decitabine activity by upregulating DCK (**Figure 1–3, S1**), we alternated THU/decitabine with THU/5-azacytidine, and examined different schedules for alternation.

**Figure 5.**
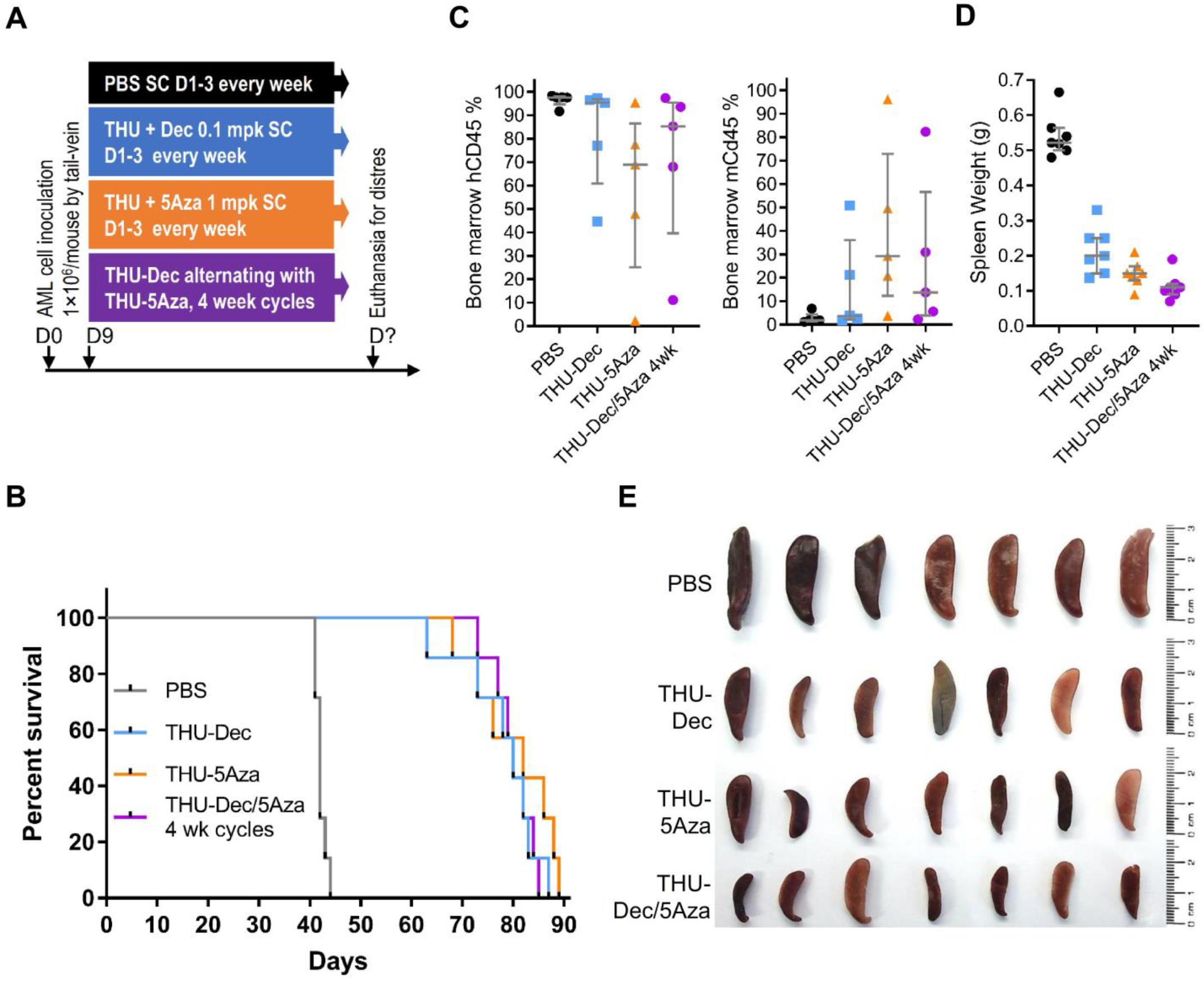
Comparison of THU/Dec alone *vs* THU/5Aza alone *vs* THU/Dec alternating with THU/5Aza in 4 week cycles. NSG mice were tail-vein inoculated with patient-derived AML cells (1×10^6^cells/mouse) and on Day 9 after inoculation randomized to the treatments as shown (n=7/group). Mice were euthanized if there were signs of distress. **A) Experiment schema**; **B) Time-to-distress and euthanasia. C) Bone marrow human (hCD45) and murine (mCd45) myelopoiesis content.** Femoral bones flushed after euthanasia. Measured by flow-cytometry. Median±IQR. **D) Spleen weights at time-of-distress/euthanasia**. Median±IQR. **E) Spleens at the time-of-distress/euthansia**.

Mice tail-vein innoculated with patient-derived AML cells (1×10^6^cells/mouse) were randomized to **(i)** vehicle; **(ii)** decitabine alone; **(iii)** THU+decitabine; or **(iv)** THU+decitabine alternating with THU+5-azacytidine week-to-week (**Figure 6A**). Median survival (time-to-distress) was impressive and greatest with THU/decitabine/5-azacytidine (221 days) *vs* THU/decitabine (180 days), decitabine-alone (111 days) or vehicle treatment (50 days) (**Figure 6B**). Stability of hemoglobin, platelet and white cell counts in mice receiving treatment every week for several months indicated a non-cytotoxic mechanism-of-action of the therapies (also previously shown^3,31,42–45^)(**Figure 6C**). There were eventual declines in hemoglobin and platelets, and increases in white cell counts (peripheral leukemia), caused by progressive leukemia (**Figure 6C**). Progressive leukemia as the cause of distress was confirmed by flow cytometry of bone marrows harvested after euthanasia which showed >95% hCD45+ cells with vehicle and 42-90% with the other treatments, with corresponding inverse amounts of murine myelopoiesis (**Figure 6D,S6**). Alternating THU/decitabine with THU/5-azacytidine in 4 week cycles, or simultaneous administration of THU/decitabine/5-azacytidine, did not add benefit over THU/decitabine alone or THU/5-azacytidine alone (**Figure 5, S5**). Thus, timing of alternation is critical.

**Figure 6.**
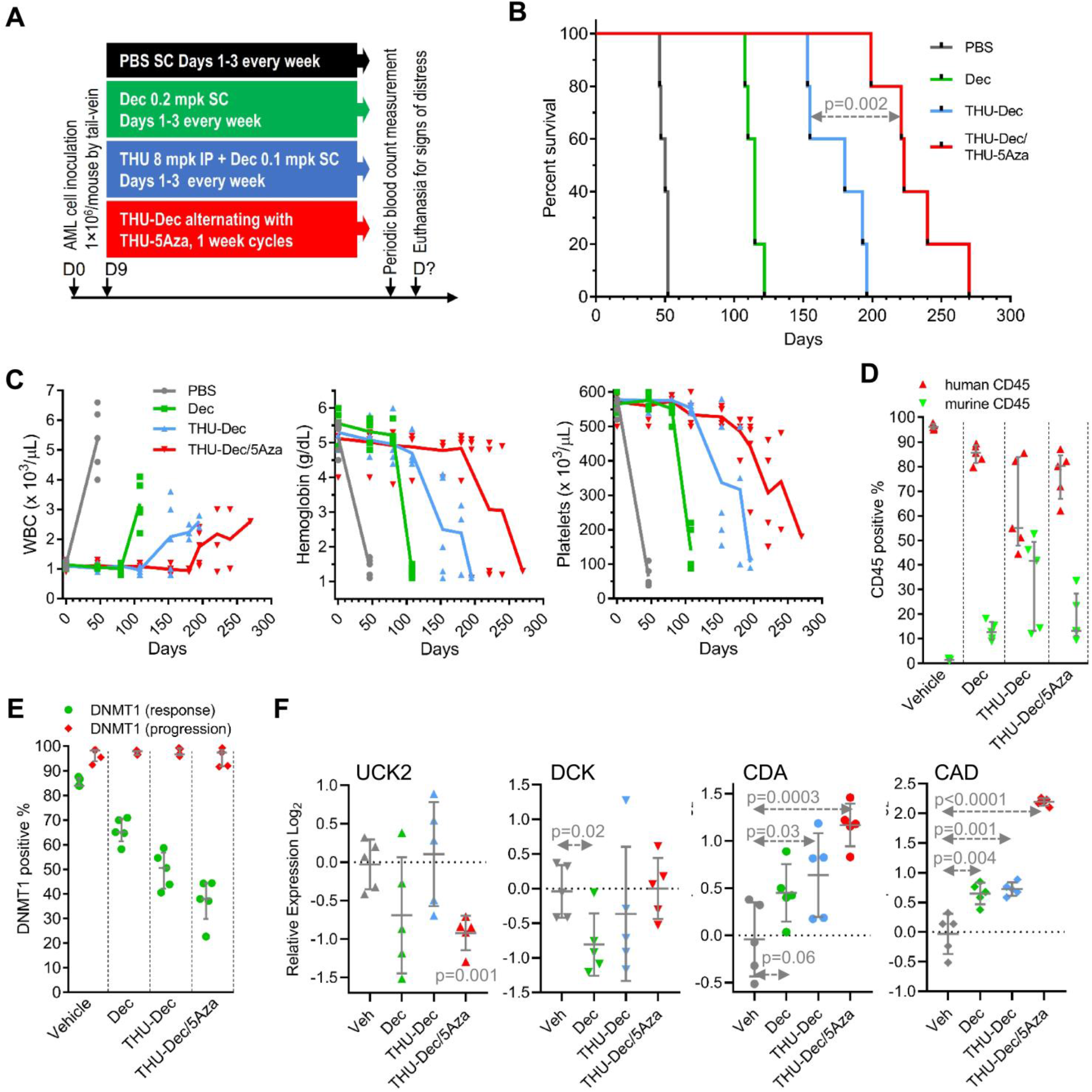
Alternating THU/decitabine with THU/5-azacytidine week to week. NSG mice were tail-vein inoculated with patient-derived AML cells (1×10^6^cells/mouse) and randomized to the treatments shown (n=5/group). Blood counts were obtained periodically by tail-vein phlebotomy. Mice were euthanized for signs of distress. **A) Experiment schema**; **B) Time-to-distress.** Log-rank test. **C) Serial blood counts**. Measured by Hemavet. Median±IQR. **D) Bone marrow replacement by AML.** Bone marrow human and murine CD45+ cells measured by flow-cytometry (**Figure S4**) after euthanasia (time-points panel B). Median±IQR. p-value Mann-Whitney test 2-sided. **E) DNMT1 was not depleted from AML cells at progression** (time-points panel B) **but was depleted at time-of-response** (bone marrow harvested at Day 63 in a separate experiment). Flow cytometry (**Figure S5**). **F) Pyrimidine metabolism gene expression in bone marrow AML cells**. QRT-PCR using human gene specific primers, bone marrow harvested after euthanasia. p-values vs vehicle, unpaired t-test, 2-sided.

### In vivo treatment-resistance

Bone marrow cells harvested at day 63 when the AML-innoculated mice were doing well on-therapy demonstrated DNMT1-depletion, with the greatest DNMT1-depletion with THU/decitabine alternating with THU/5-azacytidine week-to-week (~65% DNMT1-depletion) *vs* THU/decitabine alone (~50%) or decitabine alone (~35%) or vehicle (~15%) (**Figure 6E, S7**). By contrast, bone marrow AML cells harvested after euthanasia for distress while receiving these same therapies demonstrated preserved DNMT1 levels measured by flow-cytometry (**Figure 6E**). These in vivo treatment-resistant AML cells demonstrated significant upregulations of CDA and CAD *vs* AML cells from mice treated with vehicle, with the greatest upregulations in cells from mice that received the alternating regimen and survived the longest (**Figure 6F**). Thus, in vivo treatment-resistance was again by pyrimidine metabolism shifts adverse to DNMT1 target-engagement (**Figure S8**).

## DISCUSSION

Malignant myeloid cells proliferating through decitabine or 5-azacytidine therapy in vitro, in mice and in patients, evaded DNMT1-depletion via pyrimidine metabolism shifts adverse to pro-drug processing into the DNMT1-depleting nucleotide Aza-dCTP. Critically, the protective metabolic shifts are induced acutely - decitabine added to AML cells rapidly suppressed DCK and upregulated UCK2 and CDA, and 5-azacytidine rapidly suppressed UCK2 and upregulated DCK and CDA, with the protein expression changes peaking 72-96 hours after a single pro-drug exposure. Others have reported that decitabine upregulated CDA in leukemia and solid cancer cells by 6 to 1000-fold within 96 hours^46,47^, while 5Aza upregulated DCK in leukemia cells by ~30% within 48 hours^18^. These acute reconfigurations of pyrimidine metabolism arise from off-target actions of the agents that cause nucleotide imbalances: Decitabine inhibition of TYMS has been previously reported^38–40^, and here we found that TYMS protein is acutely depleted. A portion of administered decitabine (Aza-dC), after phosphorylation by DCK to Aza-dCMP, is deaminated by deoxycytidine deaminase (DCTD) into a deoxyuridine monophosphate (dUMP) analog Aza-dUMP. dUMP is the substrate for TYMS that rate-limits dTTP production. Aza-dUMP depletes TYMS (TYMS, like DNMT1, methylates carbon #5 of the pyrimidine ring that is substituted with a nitrogen in decitabine). In this way, decitabine decreases dTTP that in turn increases dCTP (via less dTTP inhibition of ribonucleotide reductase-mediated reduction of CDP into dCDP^38–40^). 5-azacytidine on the other hand depletes RRM1 protein and decreases dCTP^48^, presumably again as a result of the active nitrogen-substitution in the pyrimidine ring, although this has not been definitively evaluated. Stated simply, decitabine and 5-azacytidine drive dCTP levels in opposite directions, triggering distinct adaptive responses by pyrimidine metabolism, a network that senses and regulates nucleotide amounts^37^. DCK is particularly important for preserving dCTP levels, as shown by the decrease in dCTP in *DCK*-KO cells (shown also by others^49^), consistent with DCK upregulation as an appropriate adaptive metabolic reponse to dCTP suppression by 5-azacytidine. UCK2 on the other hand is particularly important for dTTP maintenance, shown by the decrease in dTTP in *UCK2*-KO cells, consistent with UCK2 upregulation as a metabolic adaptation to dTTP suppression by decitabine. Both decitabine and 5-azacytidine acutely upregulated CDA, and acutely downregulated CAD. CAD, however, was upregulated in patients’ and murine bone marrows at MDS/AML relapse/progression, the only discrepancy we found between pyrimidine metabolism patterns induced acutely versus found in stable resistant cells.

This mode of learned resistance, emerging from adaptive responses of the pyrimidine metabolism network to pro-drug perturbations, does not require mutation at the genetic level, perhaps explaining why several studies that have looked for correlations between MDS/AML genetics and decitabine/5-azacytidine resistance have generated inconclusive and even contradictory results^4,50–56^. Given that resistance-causing metabolic reconfigurations appear emergent, pre-treatment pyrimidine metabolism expression levels may also not necessarily predict response^4,50–58^. Consistent and predictable, however, were the automatic adaptive responses of pyrimidine metabolism to pro-drug exposures, enabling anticipation, outmaneuvering and even exploitation: first, serial administrations of decitabine scheduled to avoid decitabine-induced troughs in DCK expression was strikingly superior to schedules that coincided with DCK troughs, and alternating decitabine with 5-azacytidine week-to-week, timed (at least approximately) to exploit priming of each agent for activity of the other (UCK2 and DCK are maximally upregulated ~96 hrs after decitabine and 5-azacytidine respectively), was significantly superior to administration of either pro-drug alone. The timing of alternation was crucial – alternating the pro-drugs in 4 week cycles, or their simultaneous administration, did not add benefit.

Second, frequent, distributed schedules of administration, as a strategy to increase chances for overlap between malignant cell S-phase entries and drug exposure windows, were superior to conventional pulse-cycled scheduling. Such frequent, metronomic administration is feasible in mice and humans because we selected pro-drug doses to deplete DNMT1 without off-target cytotoxicity^3,4,59^. Consistent with these observations, RNA-sequencing analysis of patients’ baseline bone marrows found that a gene expression signature of low cell cycle fraction predicted non-response to standard 5-azacytidine therapy^57^, and regulatory approval of decitabine and 5-azacytidine to treat myeloid malignancies involved lowering doses from initially evaluated, toxic high doses, and the administration of these lower doses more frequently^1^

Third, combining THU, to inhibit the catabolic enzyme CDA, with decitabine and/or 5-azacytidine produced substantial extensions in anti-AML efficacy in vivo – an important detail in such combinations was that the decitabine and 5-azacytidine doses were reduced to preserve a non-cytotoxic DNMT1-targeting mode of action^3,31,44,45^. Stated another way, simple dose-escalations of decitabine or 5-azacytidine are not a solution for resistance since this compromises the therapeutic-index foundation for success: high C_max_ of these agents is cytotoxic via off-target anti-metabolite effects including depletion of TYMS (reviewed in^1^), and while AML cells that survive initial pro-drug exposures get progressively educated for resistance as they indefinitely self-replicate/proliferate, polyclonal normal myelopoiesis proliferates then differentiates in successive waves, each exposure-naïve and hence potentially vulnerable to the anti-metabolite effects of high doses of decitabine or 5-azacytidine.

The key de novo pyrimidine synthesis enzyme CAD was downregulated in AML cells upon initial challenge by the pro-drugs but was upregulated in exponentially proliferating stably resistant cells by expression and by protein S1359 phosphorylation. Although we did not find a benefit from combining decitabine with dT or hydroxyurea to inhibit ribonucleotide reductase (that is in the de novo pathway) we and others have found promise in countering resistance by inhibiting other molecular targets in the pathway, e.g., using PALA to inhibit CAD or leflunomide to inhibit DHODH^60^. Thus, in next steps, we plan to evaluate inhibitors for different targets in the de novo pyrimidine synthesis pathway, and also to increase doses of the CDA inhibitor THU, since eventual AML progression to the optimized regimen in our pre-clinical in vivo studies here was characterized by even greater upregulations of CDA.

Thus, decitabine- and 5-azacytidine-resistance emerges from adaptive responses of the pyrimidine metabolism network to the perturbations caused by these pyrimidine nucleoside analog pro-drugs. These compensatory metabolic shifts individually and collectively impede engagement of the DNMT1 molecular target of therapy. These metabolic responses, being pre-programmed and consistent, can be anticipated, outmaneuvered and even exploited, using simple and practical treatment modifications that preserve the vital therapeutic index of non-cytotoxic DNMT1-depletion.

## AUTHOR CONTRIBUTION STATEMENT

X.G., R.T., B.T., M.H., L.D., C.S., A.Z., H.C., B.H., R.S., B.K.J., E.H., J.M., Y.S. performed research and analyzed data. T.R., C.H. analyzed data. Y.S. generated hypotheses, designed research, obtained funding and wrote the paper. All authors reviewed/edited the manuscript.

## ACKNOWLEDGEMENTS

We acknowledge technical or other assistance from Quteba Ebrahem, Reda Mahfouz and Tae Hyun Hwang, and administrative support from JoAnn Bandera.

## CONFLICTS-OF-INTEREST STATEMENT

Ownership: YS – EpiDestiny. Income: none. Research support: none. Intellectual property: YS - patents around tetrahydrouridine, decitabine and 5-azacytidine (US 9,259,469 B2; US 9,265,785 B2; US 9,895,391 B2).

## SUPPLEMENTARY METHODS

### Study approvals

Bone marrow samples for research were obtained from patients with AML on a study protocol approved by the Cleveland Clinic Institutional Review Board (Cleveland, Ohio), with written informed consent prior to inclusion in the study. Experiments using patient-derived xenotransplant models of AML were approved by the Cleveland Clinic Institutional Animal Care and Use Committee (Cleveland, Ohio).

### Sources of cell lines and animals

AML cell lines OCI-AML3 were purchased from DSMZ (Braunschweig, Germany), and THP1, K562 and MOLM13 cell lines were purchased from ATCC (Manassas, Virginia). The AML cell lines, including those selected for resistance to decitabine, were authenticated (Genetica cell line testing, Burlington, NC). *DCK* and *UCK2* knock-out leukemia (HAP1) cells were engineered via Horizon Discoveries (Cambridge, United Kingdom). Primary AML cells for inoculation into NSG mice were collected with written informed consent on Cleveland Clinic Institutional Review Board approved protocol 5024. NSG mice were purchased from Jackson Laboratories (Bar Harbor, Maine).

### DNMT1 Immuno-detection and quantitation

Immunohistochemistry (IHC) was performed on decalcified and formalin-fixed paraffin embedded bone marrow biopsy sections (4μm) and on positive and negative controls (parental and DNMT1-KO HCT116 cells). Antibodies used were mouse polyclonal anti-Dnmt1 (Abcam #ab19905, Cambridge, MA), 1:200 dilution for 32 minutes at room temperature, performed with Ventana Discovery using OmniMap detection and a high pH tris-based buffer (Cell Conditioning 1, Ventana #950-124). Nuclei positive for DNMT1 were identified and quantified in high resolution, large field-of-view images per ImageIQ algorithms (Image IQ Inc., Cleveland, OH) after segmentation of images and subtraction of bone as we have previously described^4^.

DNMT1-protein measurement by flow cytometry was performed as we have previously described ^44^ using unlabeled anti-Dnmt1 antibody [EPR 3522] (0.0625 μg/test; Abcam; catalog no. ab92314) as the primary antibody

### DNA isolation, reverse transcription (RT) and real-time PCR

As we have previously described ^61^. Primer sequences were:

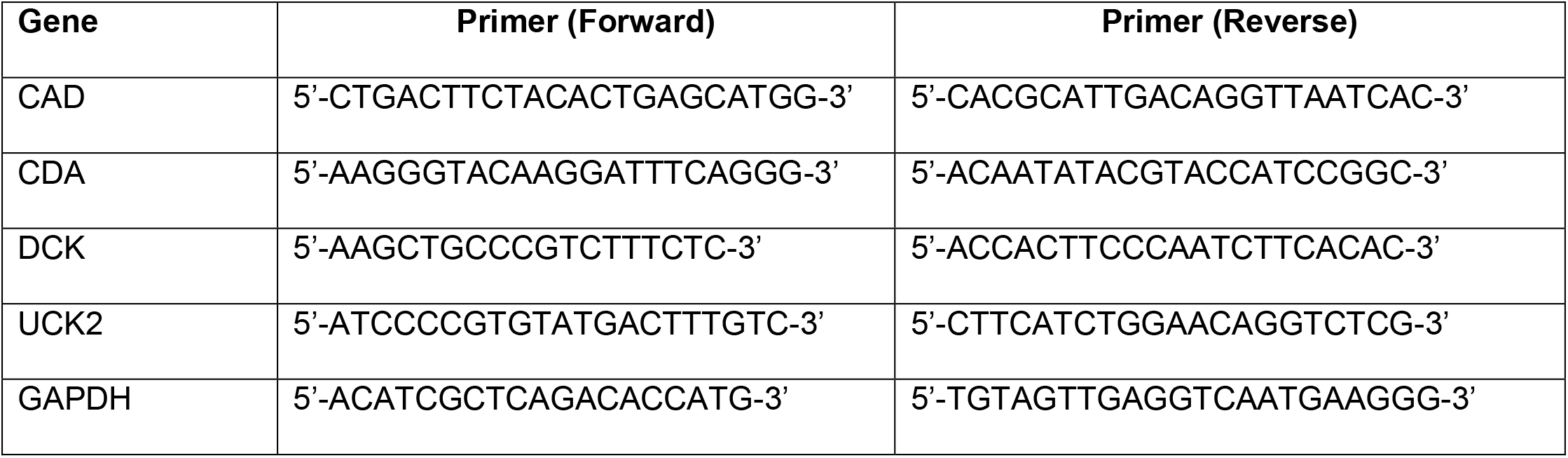

### 1D SDS-polyacrylamide gel electrophoresis and Western blot analysis

Were performed as we have previously described ^61^. Antibodies used were:

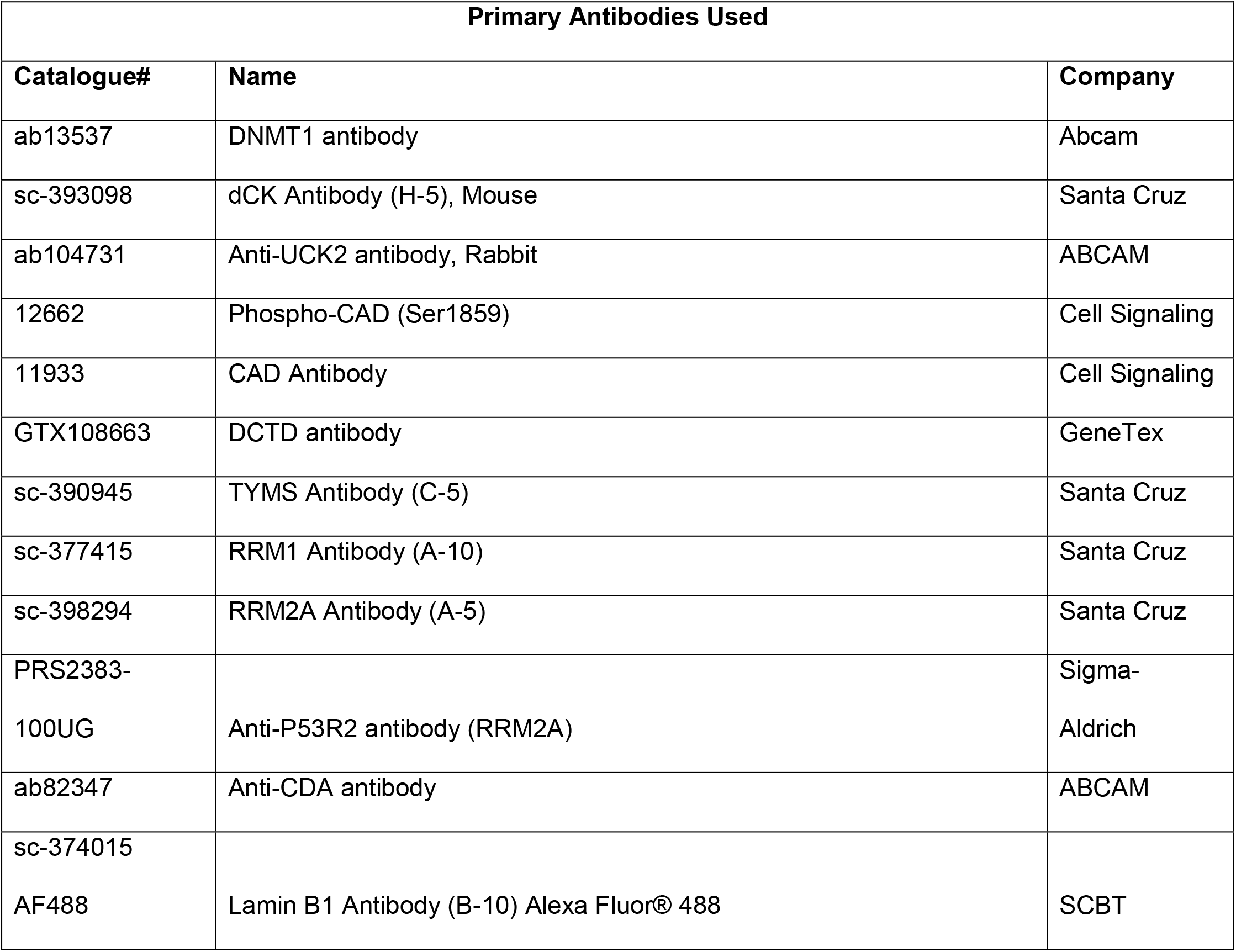
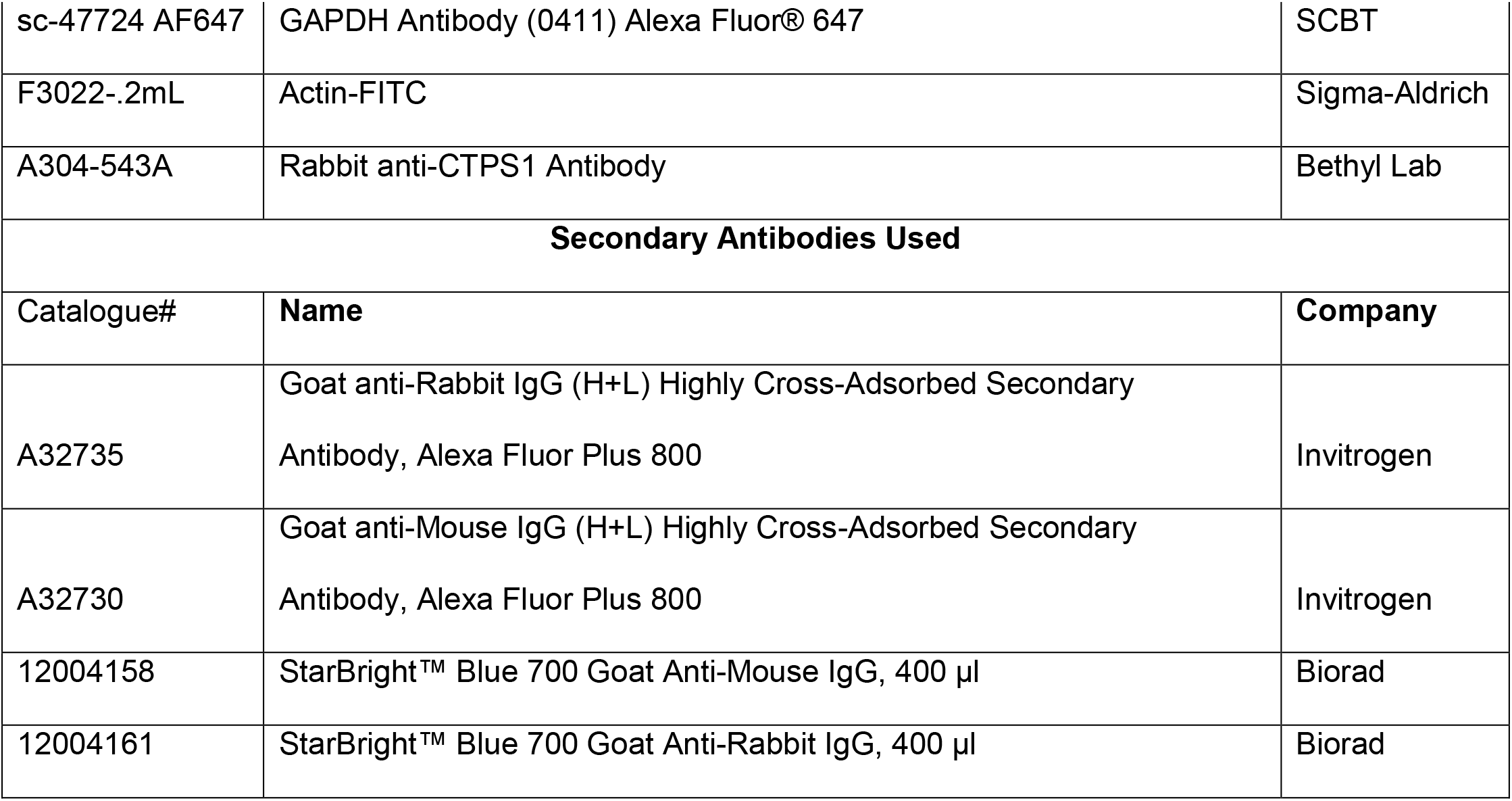

Fluorescent images were collected using Biorad’s ChemiDoc system and processed with Image lab.

### Giemsa staining of cells

As we have previously described ^61^.

### Flow Cytometry Analyses for human and murine CD45

As we have previously described ^61,62^. Antibodies used were monoclonal anti-human CD45 (clone HI30, cat. No 304016, Biolegend, 1:100) and monoclonal anti-mouse CD45 (Clone 30-F11, cat. No 1031066, Biolegend, 1:100).

### Preparation and analysis of dNTP and NTP extracts

Cells were washed twice with ice-cold 1X PBS, and counted. To lyse cells, precipitate proteins and extract nucleotides and nucleosides, for each 5-10 million cells, 250 μL of 80% acetonitrile/water was added, and the cells were incubated on ice for 15min. After incubation, the suspension was centrifuged at 140K rpm for 5min. The supernatant from this first extraction was transferred to a clean tube. The remaining pellet was extracted again with fresh 80% acetonitrile/water, and supernatant from both extractions was combined, and then evaporated to dryness using a centrifugal evaporator.

#### LCMS/MS

1mM Internal standards (13C9 15N3MP and d 13C9 15N3TP) solution (25mM ammonia acetate, 10mM DMHA, pH 8.0) was used to suspend Nucleotides and nucleosides extract. HPLC separation was carried on an ACQUITY UPLC HSS T3 Column, 100Å, 1.8 μm, 2.1 mm X 150 mm. A stepwise gradient program was applied with mobile phase A (25mM Ammonia Bicarbonate, 10mM DMHA, pH 8.0) and mobile phase B (60% Acetonitrile/water). The HPLC was interfaced with Thermofisher Quantiva triple quadruple mass spectrometer. The mass spectrometer was operated in MRM mode with optimized MRM transitions for each analyte.

#### Data analysis

Xcalibur was used to process and quantify raw data. Briefly, a processing method was built using MRM transitions and peak retention times from standards. All samples were processed with the same method to generate integrated total ion intensity (integrated peak area) for each analyte. Manual inspection was performed to confirm the peak assignment and integration. The final report value was normalized to the internal standards and total number of cells used to generate the extract.

### Treatment of a patient-derived xenotransplant model of treatment-resistant AML

Patient-derived primary AML cells from a patient with AML that had progressed on standard chemotherapy then decitabine salvage therapy, were transplanted by tail-vein injection (1.0 ×10^6^/mouse) into non-irradiated 6-8 week old NSG mice. Mice were anesthetized with isofluorane before transplantation. Mice were randomized to different treatments on Day 9 after inoculation, with treatments as indicated in each figure and legend. Doses of drugs used were: intra-peritoneal tetrahydrouridine (THU) 10 mg/kg given intra-peritoneal up to 3X/week; subcutaneous decitabine 0.2 mg/kg up to 3X/week (or 0.1 mg/kg when combined with THU); subcutaneous 5-azacytidine 2 mg/kg up to 3X/week (or 1 mg/kg when combined with THU); intra-peritoneal dT 2 g/kg up to 2X/week. Tail-vein blood samples for blood count measurement by HemaVet were obtained prior to leukemia inoculation, and at intervals thereafter as indicated in the figures. Mice were observed daily for signs of pain or distress, e.g., weight loss that exceeded 20% of initial total body weight, lethargy, vocalization, loss of motor function to any of their limbs, and were euthanized by an IACUC approved protocol if such signs were noted.

### Bioinformatic and statistical analysis

Wilcoxon rank sum, Mann Whitney, and t tests were 2-sided unless otherwise stated because of apriori literature-based hypotheses (dCTP level analyses) and performed at the 0.05 significance level or lower (Bonferroni corrections were applied for instances of multiple parallel testing). Standard deviations (SD) and inter-quartile ranges (IQR) for each set of measurements were calculated and represented as y-axis error bars on each graph. Graph Prism (GraphPad, San Diego, CA) or SAS statistical software (SAS Institute Inc., Cary, NC) was used to perform statistical analysis including correlation analyses.

---

**Figure S1.**
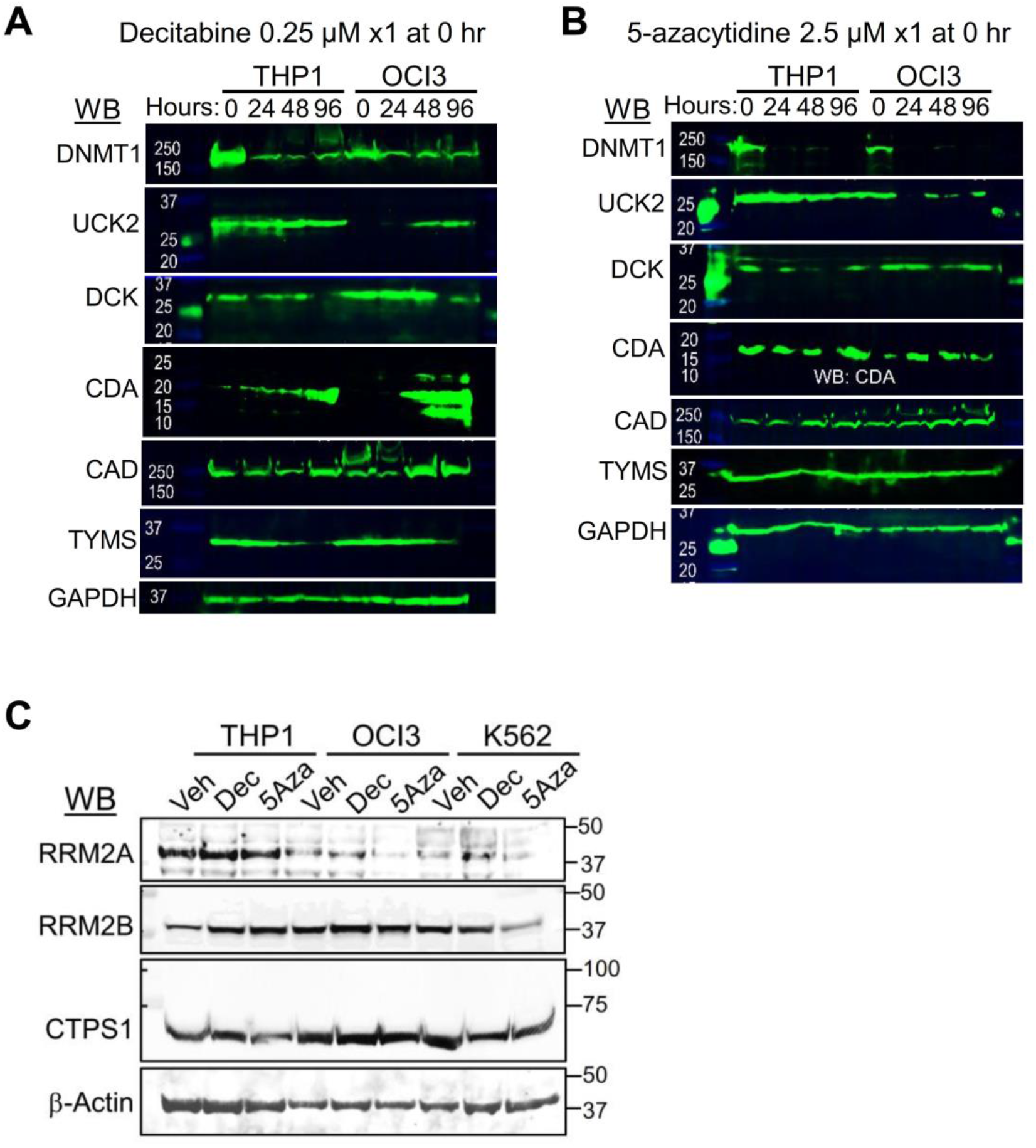
A time-course analysis suggested that peak changes in pyrimidine metabolism enzyme protein levels occurred between 48-96 hours after a single exposure to decitabine (Dec) 0.25 μM or 5-azacytidine (5Aza) 2.5 μM. **A) Western blots for DNMT1, UCK2, DCK, CDA, CAD, TYMS and GAPDH before and up to 96 hours after addition of decitabine (Dec) 0.25 μM** or **B) 5-azacytidine 2.5 μM to THP1 and OCI-AML3 cells** (added once at 0 hours). **C) 5Aza, but not Dec, decreased RRM2A levels.** Western blots 72 hours after addition of a single dose of Dec 0.25 μM or 5Aza 2.5 μM. Western blots were reproduced in biological replicates.

**Figure S2.**
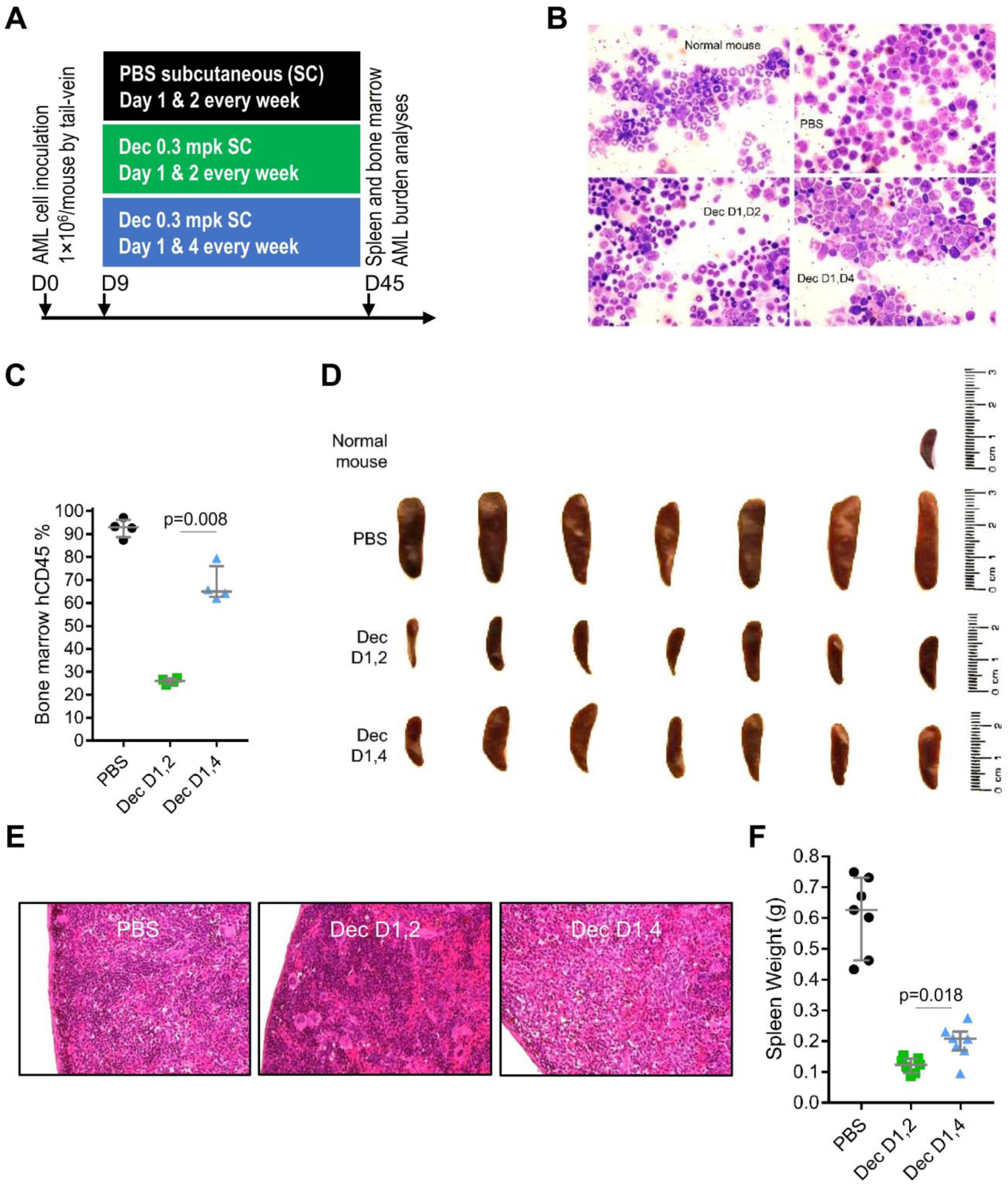
Impact of decitabine scheduling to avoid vs coincide with DCK troughs. NSG mice were tail-vein inoculated with patient-derived AML cells (1×10^6^cells/mouse) and randomized on day 9 after inoculation to (i) PBS vehicle control; (ii) subcutaneous (SC) decitabine (Dec) 0.3 mg/kg (mpk) on Day 1 and 2 of each week (D1,2); (iii) Dec 0.3 mpk on Day 1 and 4 of each week (D1,4)(n=7/group). Mice were euthanized/sacrificed on day 45 when PBS treated mice showed signs of distress. **A) Experiment schema**; **B) Bone marrow cell cytospin and Giemsa-stain at Day 45**. Normal = normal NSG mouse bone marrow; Leica DMR microscope, 630X. **C) Percentage of human CD45+ (huCD45) positive cells in bone marrow.** Flow cytometry. Median ± IQR. p-value Mann-Whitney test 2-sided. 5 mice in each treatment group analyzed. **D) Spleens at Day 45**. Normal = spleen from normal NSG mouse. **E) Spleen histology**. Hematoxylin-Eosin stain of paraffin-embedded sections. Leica DMR microscope, 400X.**F) Spleen weights.** Median±IQR. p-value Mann-Whitney test 2-sided.

**Figure S3.**
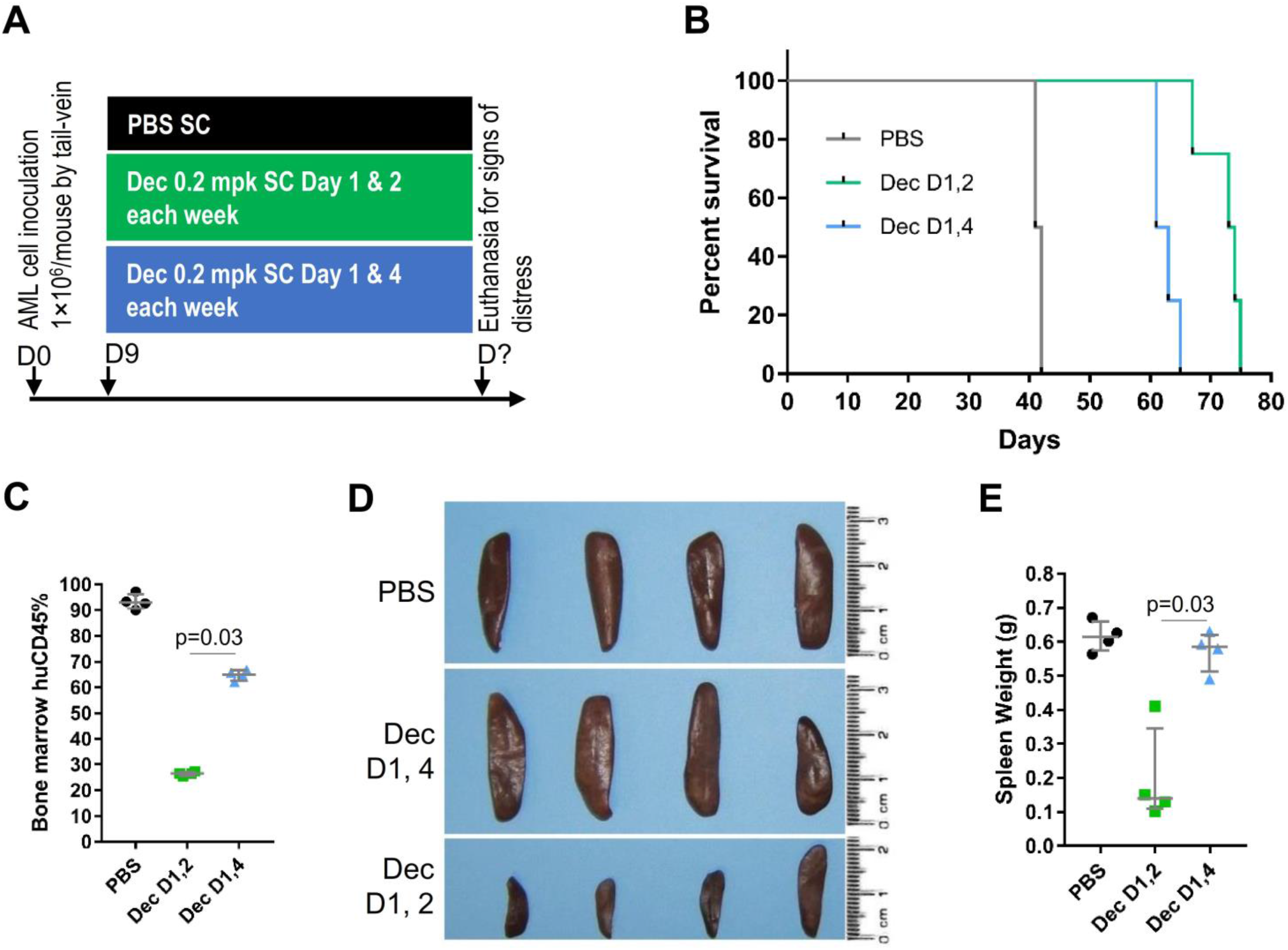
Impact of decitabine scheduling to avoid or coincide with DCK troughs. NSG mice were tail-vein inoculated with patient-derived AML cells (1×10^6^cells/mouse) and on day 9 after inoculation randomized to (i) PBS vehicle control; (ii) subcutaneous (SC) decitabine (Dec) 0.2 mg/kg (mpk) on Day 1 and 2 of each week (D1,2); (iii) Dec 0.2 mpk on Day 1 and 4 of each week (D1,4)(n=4/group). Mice were euthanized for signs of distress. **A) Experiment schema**; **B) Time-to-distress.** p-value Log-rank test. **C) Bone marrow human leukemia cell burden at time-of-distress**. Flow cytometry for human CD45+ cells. Median ± IQR. p-value Mann-Whitney test 2-sided. **D) Spleens at time-of-distress**. **E) Spleen weights at time-of-distress.** Median ± IQR. p-value Mann-Whitney test 2-sided.

**Figure S4.**
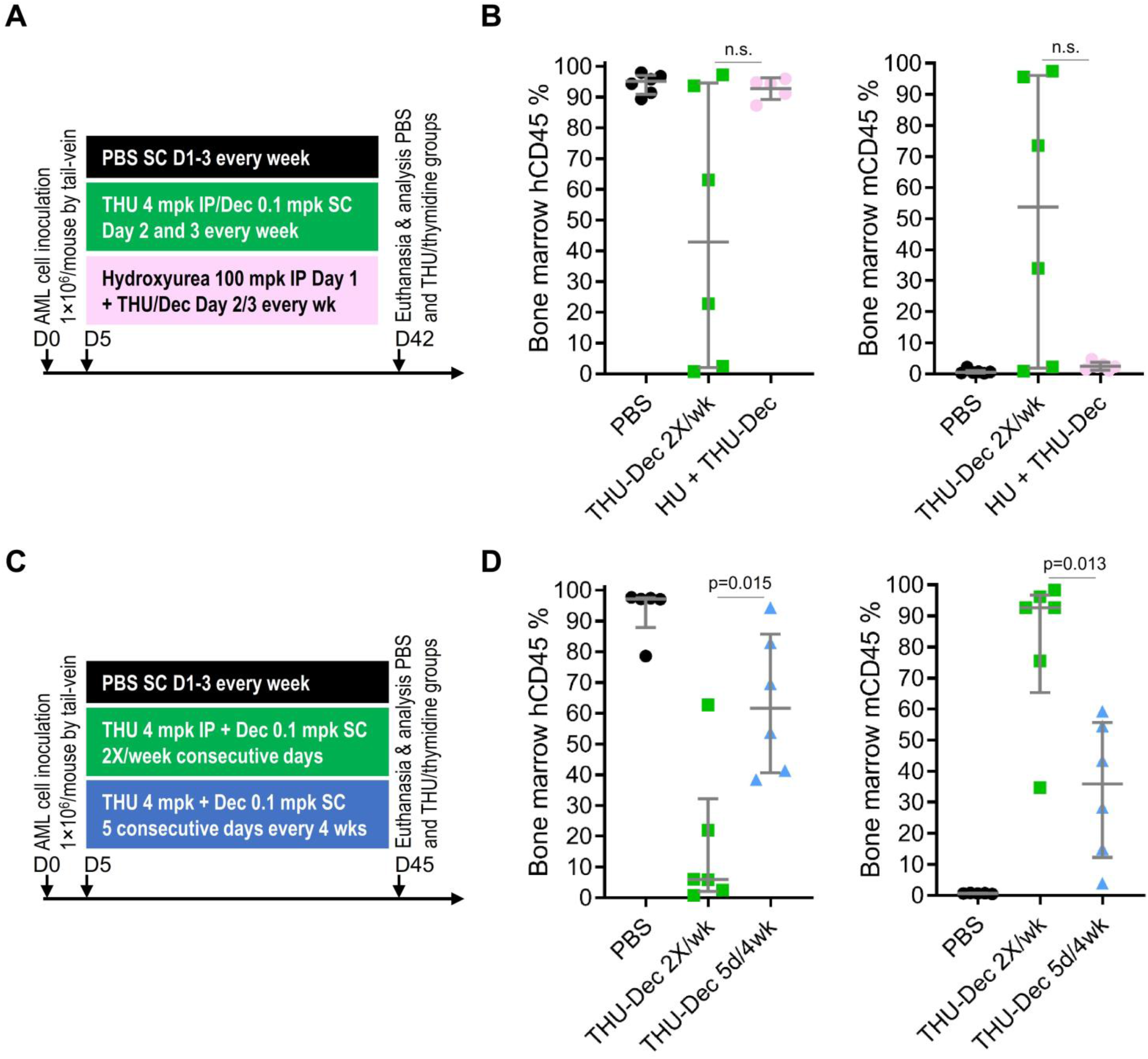
The addition of hydroxyurea to inhibit ribonucleotide reductase did not augment THU-decitabine activity; Distributed administration of THU-decitabine 2X/week was superior to pulse-cycled administration for 5 consecutive days every 4 weeks. NSG mice were tail-vein inoculated with patient-derived AML cells (1×10^6^cells/mouse) and on Day 5 after inoculation randomized to the treatments as shown (n=7/group). Mice were euthanized if there were signs of distress. **A) Experiment schema to evaluate potential benefit of adding hydroxyurea to inhibit ribonucleotide reductase**; **B) Bone marrow human (hCD45) and murine (mCd45) myelopoiesis content.** Femoral bones flushed after termination of the experiment at the time PBS-treated mice developed signs of distress. Measured by flow-cytometry. Median±IQR. P-value 2-sided Mann-Whitney test. n.s. = not significant. **C) Experiment schema to compare metronomic administration 2X/week versus pulse-cycled administration for 5 consecutive days every 4 weeks**; **D) Bone marrow human (hCD45) and murine (mCd45) myelopoiesis content.** Femoral bones flushed after termination of the experiment at the time PBS-treated mice developed signs of distress. Measured by flow-cytometry. Median±IQR. P-value 2-sided Mann-Whitney test. n.s. = not significant.

**Figure S5.**
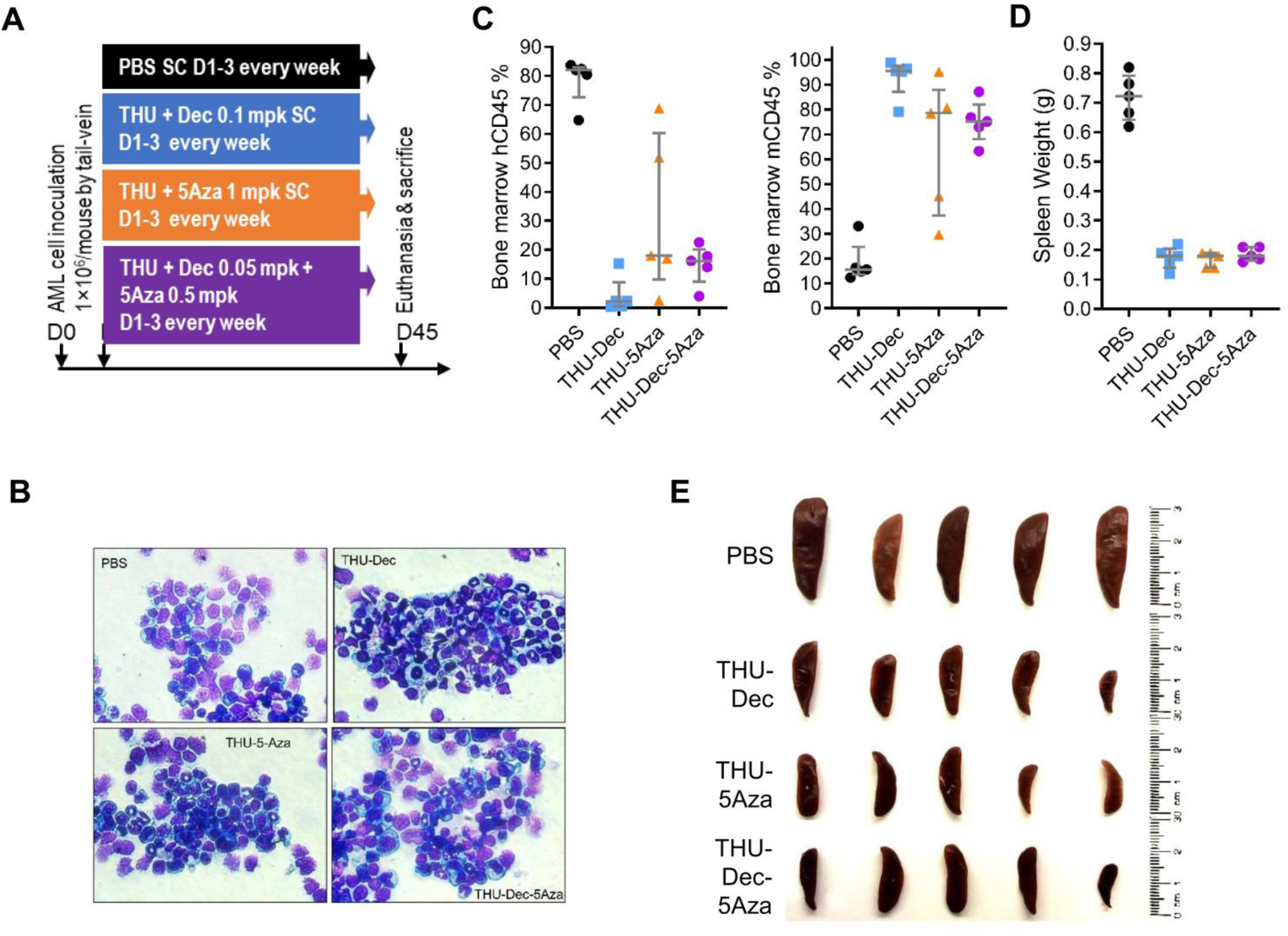
THU/decitabine *vs* THU/5-azacytidine *vs* THU/decitabine/5-azacytidine. NSG mice were tail-vein inoculated with patient-derived AML cells (1×10^6^cells/mouse) and on Day 9 after inoculation randomized to the treatments as shown (n=5/group). All mice were euthanized or sacrificed when the vehicle-treated group became distressed at Day 45. **A) Experiment schema**; **B) Giemsa stained cytospins of bone marrow cells.** Flushed from femoral bones after euthanasia. Magnification 630X. Leica DMR microscope.. **C) Bone marrow human (hCD45) and murine (mCd45) myelopoiesis content.** Measured by flow-cytometry. Median±IQR. **D) Spleen weights at time-of-distress/euthanasia**. Median±IQR. **E) Spleens**.

**Figure S6.**
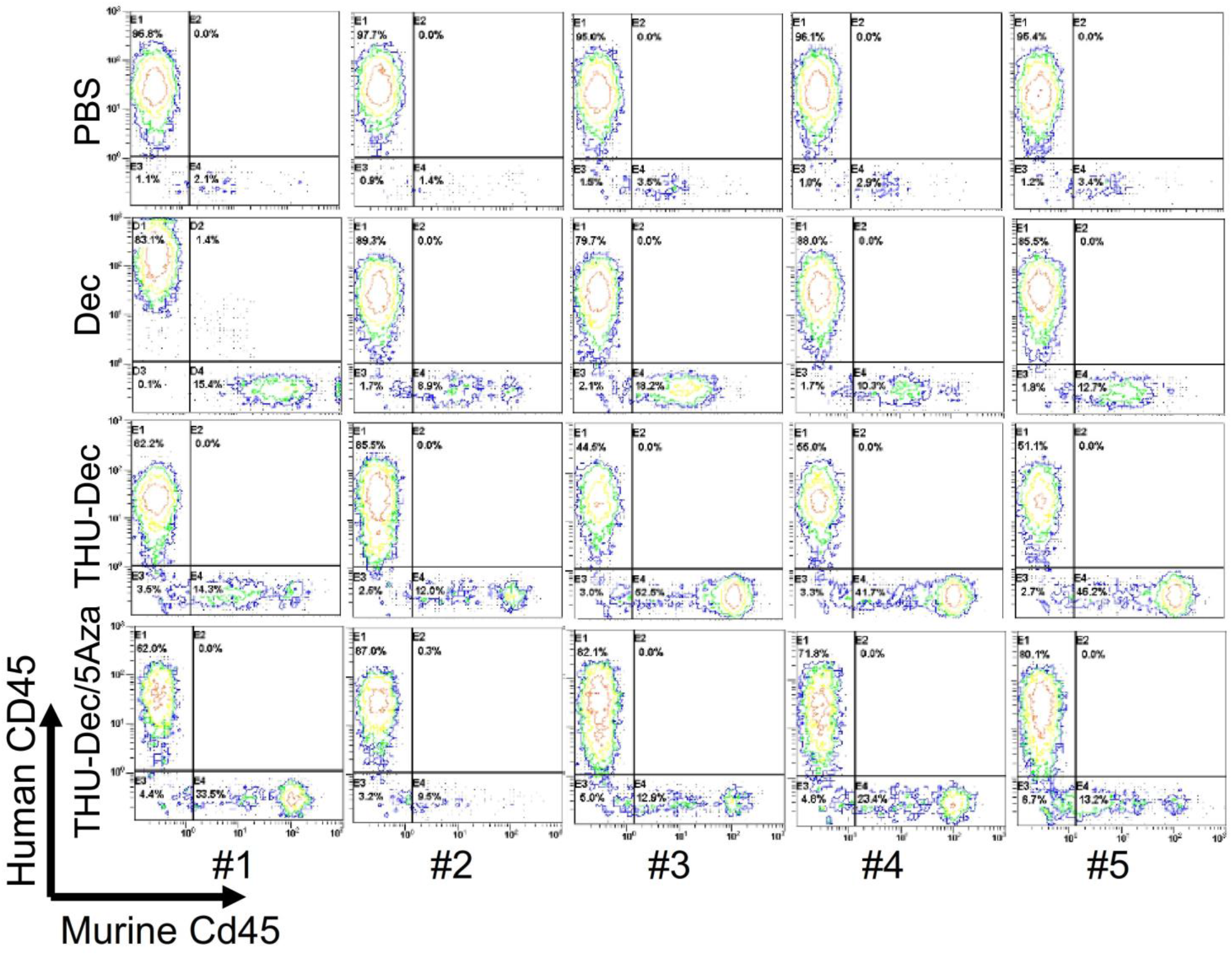
Bone marrow AML burden (human CD45) versus murine myelopoiesis (murine Cd45) in bone marrow cells harvested after euthanasia for distress (time-points as per Figure 7B).

**Figure S7.**
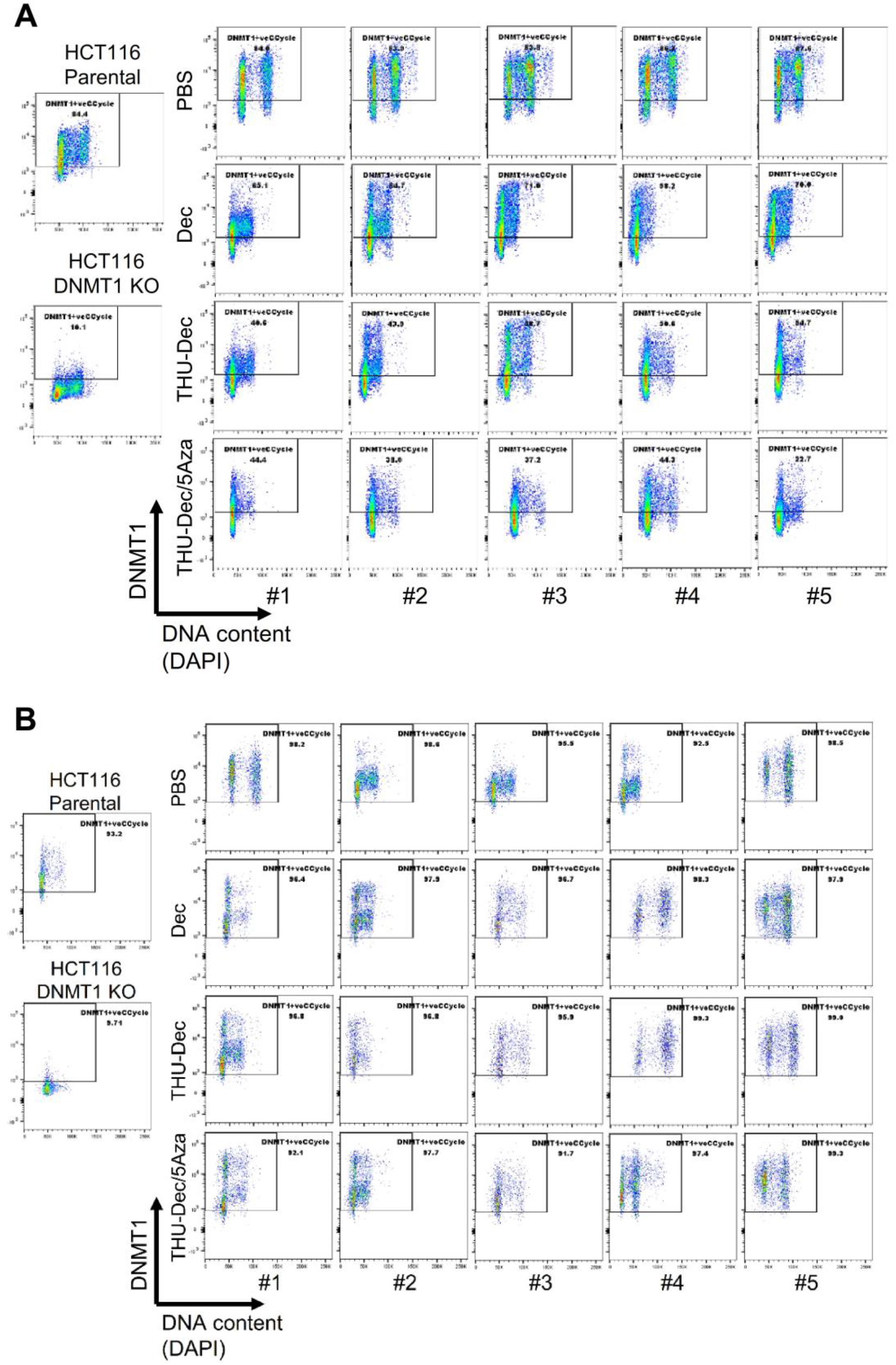
DNMT1 was not depleted from AML cells at progression (time-of-distress as shown in Figure 7B) but was depleted at time-of-response. (bone marrow harvested at Day 63 in a separate experiment). Flow cytometry. Positive and negative controls for DNMT1 were HCT116 colon cancer cells with wild-type DNMT1 and DNMT1 knock-out (KO).

**Figure S8.**
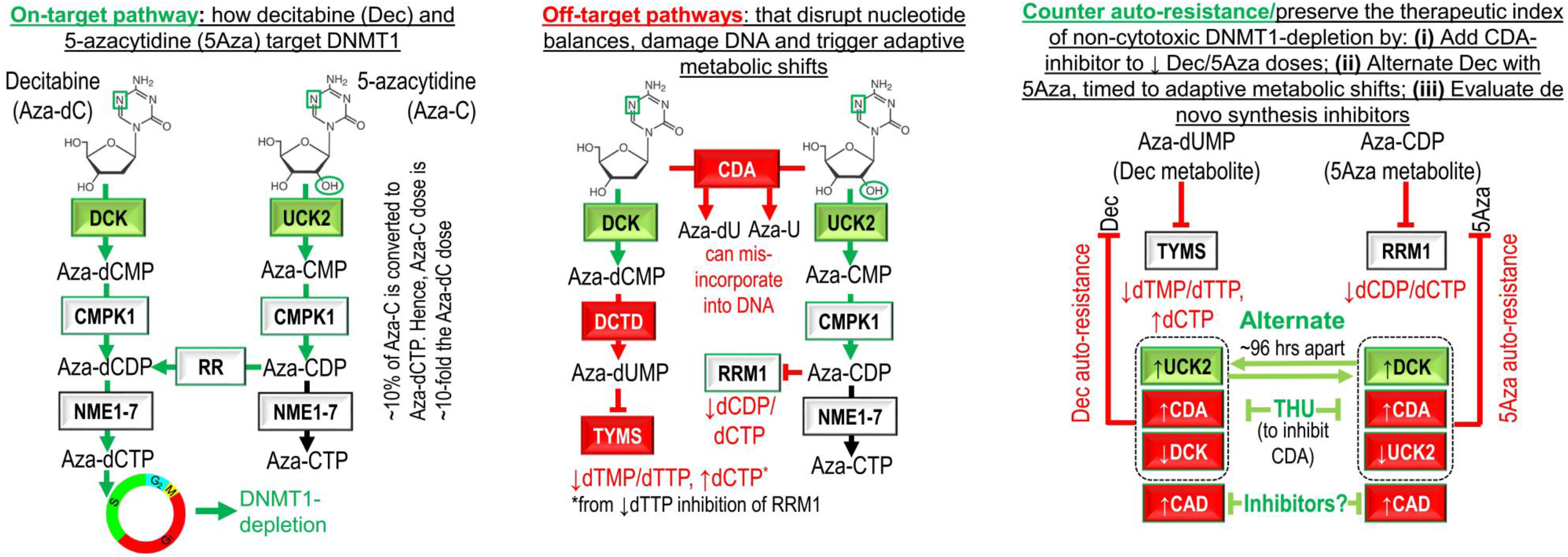
Summary of on-target and off-target pathways, and potential solutions for resistance.

## Notes

**Financial support:** National Institutes of Health P30 CA043703

## REFERENCES

1. Saunthararajah Y. Key clinical observations after 5-azacytidine and decitabine treatment of myelodysplastic syndromes suggest practical solutions for better outcomes. Hematology Am Soc Hematol Educ Program. 2013;2013:511–521.

2. Vesely J, Cihak A, Sorm F. Characteristics of mouse leukemic cells resistant to 5-azacytidine and 5-aza-2’-deoxycytidine. Cancer Res. 1968;28(10):1995–2000.

3. Ng KP, Ebrahem Q, Negrotto S, et al. p53 independent epigenetic-differentiation treatment in xenotransplant models of acute myeloid leukemia. Leukemia. 2011;25(11):1739–1750.

4. Saunthararajah Y, Sekeres M, Advani A, et al. Evaluation of noncytotoxic DNMT1-depleting therapy in patients with myelodysplastic syndromes. J Clin Invest. 2015;125(3):1043–1055.

5. Tsai HC, Li H, Van Neste L, et al. Transient Low Doses of DNA-Demethylating Agents Exert Durable Antitumor Effects on Hematological and Epithelial Tumor Cells. Cancer Cell. 2012;21(3):430–446.

6. Negrotto S, Ng KP, Jankowska AM, et al. CpG methylation patterns and decitabine treatment response in acute myeloid leukemia cells and normal hematopoietic precursors. Leukemia. 2012;26(2):244–254.

7. Momparler RL, Cote S, Momparler LF. Epigenetic action of decitabine (5-aza-2’-deoxycytidine) is more effective against acute myeloid leukemia than cytotoxic action of cytarabine (ARA-C). Leuk Res. 2013.

8. Trowbridge JJ, Sinha AU, Zhu N, Li M, Armstrong SA, Orkin SH. Haploinsufficiency of Dnmt1 impairs leukemia stem cell function through derepression of bivalent chromatin domains. Genes Dev. 2012;26(4):344–349.

9. Broske AM, Vockentanz L, Kharazi S, et al. DNA methylation protects hematopoietic stem cell multipotency from myeloerythroid restriction. NatGenet. 2009;41(11):1207–1215.

10. Milhem M, Mahmud N, Lavelle D, et al. Modification of hematopoietic stem cell fate by 5aza 2 ’ deoxycytidine and trichostatin A. Blood. 2004;103(11):4102–4110.

11. Hu Z, Negrotto S, Gu X, et al. Decitabine maintains hematopoietic precursor self-renewal by preventing repression of stem cell genes by a differentiation-inducing stimulus. Mol Cancer Ther. 2010;9(6):1536–1543.

12. Chaurasia P, Gajzer DC, Schaniel C, D’Souza S, Hoffman R. Epigenetic reprogramming induces the expansion of cord blood stem cells. J Clin Invest. 2014.

13. Araki H, Yoshinaga K, Boccuni P, Zhao Y, Hoffman R, Mahmud N. Chromatin-modifying agents permit human hematopoietic stem cells to undergo multiple cell divisions while retaining their repopulating potential. Blood. 2007;109(8):3570–3578.

14. Suzuki M, Harashima A, Okochi A, et al. 5-Azacytidine supports the long-term repopulating activity of cord blood CD34(+) cells. AmJHematol. 2004;77(3):313–315.

15. Velcheti V, Schrump D, Saunthararajah Y. Ultimate Precision: Targeting Cancer but Not Normal Self-replication. Am Soc Clin Oncol Educ Book. 2018(38):950–963.

16. Stegmann AP, Honders WH, Willemze R, Ruiz vHV, Landegent JE. Transfection of wild-type deoxycytidine kinase (dck) cDNA into an AraC- and DAC-resistant rat leukemic cell line of clonal origin fully restores drug sensitivity. Blood. 1995;85(5):1188–1194.

17. Wang H, Chen P, Wang J, et al. In vivo quantification of active decitabine-triphosphate metabolite: a novel pharmacoanalytical endpoint for optimization of hypomethylating therapy in acute myeloid leukemia. AAPS J. 2013;15(1):242–249.

18. Antonsson BE, Avramis VI, Nyce J, Holcenberg JS. Effect of 5-azacytidine and congeners on DNA methylation and expression of deoxycytidine kinase in the human lymphoid cell lines CCRF/CEM/0 and CCRF/CEM/dCk-1. Cancer Res. 1987;47(14):3672–3678.

19. Qin T, Jelinek J, Si J, Shu J, Issa JP. Mechanisms of resistance to 5-aza-2’-deoxycytidine in human cancer cell lines. Blood. 2009;113(3):659–667.

20. Grant S, Bhalla K, Gleyzer M. Effect of uridine on response of 5-azacytidine-resistant human leukemic cells to inhibitors of de novo pyrimidine synthesis. Cancer Res. 1984;44(12 Pt 1):5505–5510.

21. Liacouras AS, Anderson EP. Uridine-cytidine kinase. IV. Kinetics of the competition between 5-azacytidine and the two natural substrates. Mol Pharmacol. 1979;15(2):331–340.

22. Sripayap P, Nagai T, Uesawa M, et al. Mechanisms of resistance to azacitidine in human leukemia cell lines. Exp Hematol. 2014;42(4):294–306 e292.

23. Qin T, Castoro R, El Ahdab S, et al. Mechanisms of resistance to decitabine in the myelodysplastic syndrome. PLoS One. 2011;6(8):e23372.

24. Wu P, Geng S, Weng J, et al. The hENT1 and DCK genes underlie the decitabine response in patients with myelodysplastic syndrome. Leuk Res. 2015;39(2):216–220.

25. Camiener GW, Smith CG. Studies of the enzymatic deamination of cytosine arabinoside. I. Enzyme distribution and species specificity. BiochemPharmacol. 1965;14(10):1405–1416.

26. Zauri M, Berridge G, Thezenas ML, et al. CDA directs metabolism of epigenetic nucleosides revealing a therapeutic window in cancer. Nature. 2015;524(7563):114–118.

27. Beausejour CM, Eliopoulos N, Momparler L, Le NL, Momparler RL. Selection of drug-resistant transduced cells with cytosine nucleoside analogs using the human cytidine deaminase gene. Cancer Gene Ther. 2001;8(9):669–676.

28. Eliopoulos N, Cournoyer D, Momparler RL. Drug resistance to 5-aza-2’-deoxycytidine, 2’,2’-difluorodeoxycytidine, and cytosine arabinoside conferred by retroviral-mediated transfer of human cytidine deaminase cDNA into murine cells. Cancer ChemotherPharmacol. 1998;42(5):373–378.

29. Liu Z, Marcucci G, Byrd JC, Grever M, Xiao J, Chan KK. Characterization of decomposition products and preclinical and low dose clinical pharmacokinetics of decitabine (5-aza-2’-deoxycytidine) by a new liquid chromatography/tandem mass spectrometry quantification method. Rapid CommunMass Spectrom. 2006;20(7):1117–1126.

30. Liu Z, Liu S, Xie Z, et al. Characterization of in vitro and in vivo hypomethylating effects of decitabine in acute myeloid leukemia by a rapid, specific and sensitive LC-MS/MS method. Nucleic Acids Res. 2007;35(5):e31.

31. Ebrahem Q, Mahfouz R, Ng KP, Saunthararajah Y. High cytidine deaminase expression in the liver provides sanctuary for cancer cells from decitabine treatment effects. Oncotarget. 2012;3(10):1137–1145.

32. Mahfouz RZ, Jankowska A, Ebrahem Q, et al. Increased CDA expression/activity in males contributes to decreased cytidine analog half-life and likely contributes to worse outcomes with 5-azacytidine or decitabine therapy. Clin Cancer Res. 2013;19(4):938–948.

33. Mahfouz RZ, Koh LS, Teo M, Chee CL, Toh HC, Saunthararajah Y. Gender, cytidine deaminase, and 5-aza/decitabine--response. Clin Cancer Res. 2013;19(11):3106–3107.

34. DeZern AE, Zeidan AM, Barnard J, et al. Differential response to hypomethylating agents based on sex: a report on behalf of the MDS Clinical Research Consortium (MDS CRC). Leuk Lymphoma. 2017;58(6):1325–1331.

35. Grant S, Bhalla K, Gleyzer M. Interaction of deoxycytidine and deoxycytidine analogs in normal and leukemic human myeloid progenitor cells. LeukRes. 1986;10(9):1139–1146.

36. Ng SK, Rogers J, Sanwal BD. Alterations in differentiation and pyrimidine pathway enzymes in 5-azacytidine resistant variants of a myoblast line. J Cell Physiol. 1977;90(2):361–347.

37. Lane AN, Fan TW. Regulation of mammalian nucleotide metabolism and biosynthesis. Nucleic Acids Res. 2015;43(4):2466–2485.

38. Almqvist H, Axelsson H, Jafari R, et al. CETSA screening identifies known and novel thymidylate synthase inhibitors and slow intracellular activation of 5-fluorouracil. Nat Commun. 2016;7:11040.

39. Heinemann V, Plunkett W. Modulation of deoxynucleotide metabolism by the deoxycytidylate deaminase inhibitor 3,4,5,6-tetrahydrodeoxyuridine. Biochem Pharmacol. 1989;38(22):4115–4121.

40. Bianchi V, Pontis E, Reichard P. Regulation of pyrimidine deoxyribonucleotide metabolism by substrate cycles in dCMP deaminase-deficient V79 hamster cells. Mol Cell Biol. 1987;7(12):4218–4224.

41. Nathanson DA, Armijo AL, Tom M, et al. Co-targeting of convergent nucleotide biosynthetic pathways for leukemia eradication. J Exp Med. 2014;211(3):473–486.

42. Gu X, Ebrahem Q, Mahfouz RZ, et al. Leukemogenic nucleophosmin mutation disrupts the transcription factor hub that regulates granulomonocytic fates. J Clin Invest. 2018;128(10):4260–4279.

43. Saunthararajah Y, Hillery CA, Lavelle D, et al. Effects of 5-aza-2 ’-deoxycytidine on fetal hemoglobin levels, red cell adhesion, and hematopoietic differentiation in patients with sickle cell disease. Blood. 2003;102(12):3865–3870.

44. Molokie R, Lavelle D, Gowhari M, et al. Oral tetrahydrouridine and decitabine for non-cytotoxic epigenetic gene regulation in sickle cell disease: A randomized phase 1 study. PLoS Med. 2017;14(9):e1002382.

45. Lavelle D, Vaitkus K, Ling Y, et al. Effects of tetrahydrouridine on pharmacokinetics and pharmacodynamics of oral decitabine. Blood. 2012;119(5):1240–1247.

46. Momparler RL, Laliberte J. Induction of cytidine deaminase in HL-60 myeloid leukemic cells by 5-aza-2’-deoxycytidine. LeukRes. 1990;14(9):751–754.

47. Mameri H, Bieche I, Meseure D, et al. Cytidine Deaminase Deficiency Reveals New Therapeutic Opportunities against Cancer. Clin Cancer Res. 2017;23(8):2116–2126.

48. Aimiuwu J, Wang H, Chen P, et al. RNA-dependent inhibition of ribonucleotide reductase is a major pathway for 5-azacytidine activity in acute myeloid leukemia. Blood. 2012;119(22):5229–5238.

49. Austin WR, Armijo AL, Campbell DO, et al. Nucleoside salvage pathway kinases regulate hematopoiesis by linking nucleotide metabolism with replication stress. J Exp Med. 2012;209(12):2215–2228.

50. Im AP, Sehgal AR, Carroll MP, et al. DNMT3A and IDH mutations in acute myeloid leukemia and other myeloid malignancies: associations with prognosis and potential treatment strategies. Leukemia. 2014;28(9):1774–1783.

51. Voso MT, Santini V, Fabiani E, et al. Why methylation is not a marker predictive of response to hypomethylating agents. Haematologica. 2014;99(4):613–619.

52. DiNardo CD, Patel KP, Garcia-Manero G, et al. Lack of association of IDH1, IDH2 and DNMT3A mutations with outcome in older patients with acute myeloid leukemia treated with hypomethylating agents. Leuk Lymphoma. 2014;55(8):1925–1929.

53. Traina F, Visconte V, Elson P, et al. Impact of molecular mutations on treatment response to DNMT inhibitors in myelodysplasia and related neoplasms. Leukemia. 2014;28(1):78–87.

54. Metzeler KH, Walker A, Geyer S, et al. DNMT3A mutations and response to the hypomethylating agent decitabine in acute myeloid leukemia. Leukemia. 2012;26(5):1106–1107.

55. Bejar R, Lord A, Stevenson K, et al. TET2 mutations predict response to hypomethylating agents in myelodysplastic syndrome patients. Blood. 2014;124(17):2705–2712.

56. Braun T, Itzykson R, Renneville A, et al. Molecular predictors of response to decitabine in advanced chronic myelomonocytic leukemia: a phase 2 trial. Blood. 2011;118(14):3824–3831.

57. Unnikrishnan A, Papaemmanuil E, Beck D, et al. Integrative Genomics Identifies the Molecular Basis of Resistance to Azacitidine Therapy in Myelodysplastic Syndromes. Cell Rep. 2017;20(3):572–585.

58. Valencia A, Masala E, Rossi A, et al. Expression of nucleoside-metabolizing enzymes in myelodysplastic syndromes and modulation of response to azacitidine. Leukemia. 2014;28(3):621–628.

59. Awada H, Mahfouz RZ, Kishtagari A, et al. Extended experience with a non-cytotoxic DNMT1-targeting regimen of decitabine to treat myeloid malignancies. Br J Haematol. 2019.

60. Imanishi S, Takahashi R, Katagiri S, et al. Teriflunomide restores 5-azacytidine sensitivity via activation of pyrimidine salvage in 5-azacytidine-resistant leukemia cells. Oncotarget. 2017;8(41):69906–69915.

61. Gu X, Hu Z, Ebrahem Q, et al. Runx1 regulation of Pu.1 corepressor/coactivator exchange identifies specific molecular targets for leukemia differentiation therapy. J Biol Chem. 2014;289(21):14881–14895.

62. Hu Z, Gu X, Baraoidan K, et al. RUNX1 regulates corepressor interactions of PU.1. Blood. 2011;117(24):6498–6508.

